# Regulatory architecture of disease resistance in maize revealed by multi-omic systems genetics

**DOI:** 10.1101/2024.08.29.610401

**Authors:** Natalie M Clark, Gaoyuan Song, Mercy K. Kabahuma, Judith M Kolkman, Shawn A Christensen, Christian Montes, Shikha Malik, Rebecca J Nelson, Justin W Walley

## Abstract

Complex traits such as disease resistance have been traditionally studied using quantitative genetics. Here, we use systems genetics to integrate disease severity and multi-omic quantitate trait loci (QTL) to uncover biological networks underlying interaction with northern leaf blight (NLB), a yield-limiting disease of corn. Specifically, we integrated transcriptome, (phospho)proteome, and metabolome measurements to map molecular QTL and build predictive regulatory networks following NLB infection. These inferred networks identified a critical signaling module that was genetically validated comprised of a kinase termed <u>N</u>LB <u>S</u>USCEPTIBLE <u>K</u>INASE 1, a bHLH transcription factor, and the lignin biosynthesis enzyme BROWN MIDRIB 2. Our results demonstrate the feasibility of high-throughput mapping of genetic determinants of gene-product levels and demonstrates the power of systems genetics to identify upstream regulatory genes that confer resistance to NLB that can inform future strategies for crop protection.

## Introduction

Northern leaf blight (NLB), caused by the fungal pathogen *Setosphaeria turcica* (anamorph: *Exserohilum turcica*) is one of the most important foliar diseases affecting maize production worldwide, resulting in severe yield losses (*1*). Genetic resistance to NLB is comprised of both major qualitative resistance genes and quantitative trait loci (QTLs). The use of major resistance genes remains elusive as breeding targets due to racial structure of endogenous pathogen isolates, and the complexity of the multiple allele HtN1/Ht2/Ht3 gene complex. Characterization and of QTLs, i.e. minor resistance genes, for disease resistance have been described for NLB with several genes conferring minor resistance described, yet we lack a comprehensive understanding of NLB immune signaling (*2–6*).

While major strides have been made to dissect the genomic variations affecting maize diseases such as NLB, these studies do not currently incorporate the impact of post-translational modifications (PTMs). Importantly, PTMs affect protein function in many ways including altering protein stability, activity, and/or subcellular localization (*7*). Protein phosphorylation is a prevalent post-translational modification that is intricately involved in many signaling pathways and biological processes including being an integral component of biotic stress signaling (*8*, *9*).

Based on the development of high throughput sequencing methods and mass spectrometry (MS) based proteomics technologies, transcript abundance (eQTL) or protein abundance QTL (pQTL) have been mapped for different organisms to understand the regulatory mechanisms behind complex traits (*10–14*). Recent technological improvements in high-resolution MS and quantitative phosphoproteomics provide the potential for genome-wide phosphorylation site QTL (phosQTL) mapping (*15*, *16*). However, there are still some challenges to performing these at scale as well as to connect genomic variations, protein phosphorylation, and quantitative traits such as disease resistance.

To characterize the regulatory mechanisms underlying the defense response against NLB and to investigate novel candidate genes and signaling pathways which may confer NLB resistance, we performed a multi-QTL experiment including disease phenotype (dQTL), eQTL, pQTL, phosQTL, and metabolite QTL (mQTL) with NLB treated Intermated B73xMo17 Doubled Haploid Lines (IBMDHLs). We constructed an integrative regulatory network with the QTL mapped molecular traits (transcripts, proteins, phosphorylation sites, and metabolites). This combinatorial multi-QTL mapping and network inference approach identified a novel signaling pathway between a kinase we term NLB SUSCEPTIBLE KINASE 1 (NSK1; Zm00001d037297), the basic helix-loop-helix transcription factor bHLH106, and the lignin biosynthesis enzyme BROWN MIDRIB 2 (BM2). This molecular pathway was functionally validated by follow-up experiments that link BM2 expression to both NSK1 and bHLH106. Critically, the loss of either NSK1 or bHLH106 in CRISPR-induced mutant maize lines increases susceptibility to NLB. This study illustrates the power of multi-omic based QTL mapping approaches in identifying novel signaling modules regulating complex biological processes such as disease resistance.

### Multi-omics QTL mapping

To generate a multi-omics profile of NLB-infected maize leaves, we planted 134 Intermated B73xMo17 Doubled Haploid Lines (IBMDHLs) along with the B73 and Mo17 parental lines in an agricultural research field outside of Boone, IA in the summer of 2018 (Supplemental Figure 1). Leaves of V6-V7 maize plants were inoculated with NLB and leaf tissue was collected 7 days after infection. We then performed 3’ QuantSeq (*17*) to measure transcriptome levels and measured protein abundance and phosphorylation state using two-dimensional liquid chromatography-tandem mass spectrometry (2D-LC-MS/MS) on Tandem Mass Tag (TMT) labeled peptides (*18*, *19*) (Supplemental Figure 1). Using these methods, we quantified 34,816 transcripts, 11,618 protein groups, and 42,078 phosphosites (Figure 1a, Supplemental Datasets 1-3).

**Figure 1.**
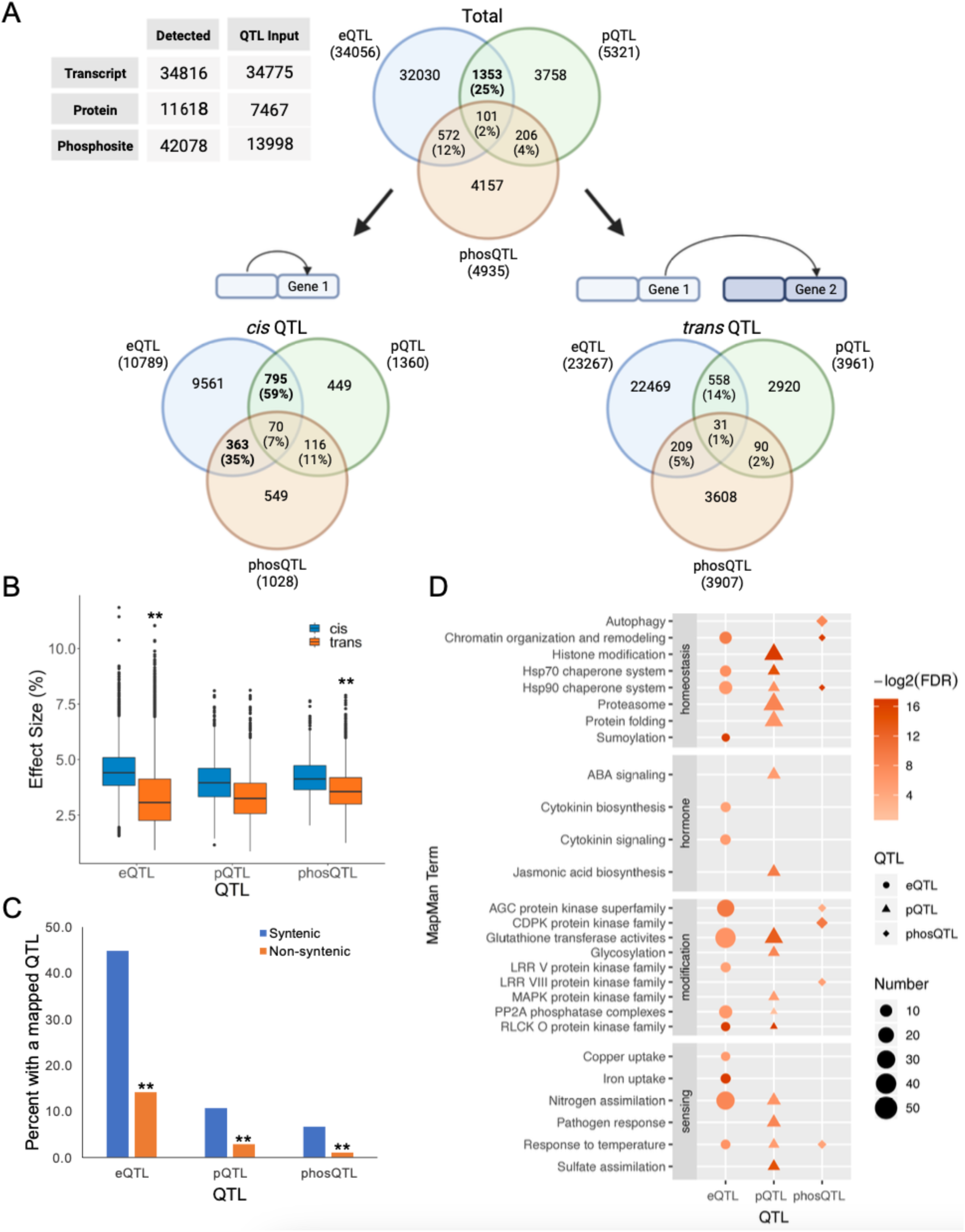
Multi-omics QTL mapping of NLB-infected maize. a) (left, table) Detected gene products. Only gene products detected in at least half (67) of the 134 lines were used for QTL mapping. (right, Venn Diagrams) Overlap of eQTL, pQTL, and phosQTL separated by *cis* and *trans* QTL. Bold numbers are statistically significant, *p* < 0.0001, hypergeometric test. (b) Effect size of eQTL, pQTL, and phosQTL separated by *cis* (blue) and *trans* (orange) QTL. ** denotes *p* < 0.0001, Wilcoxon rank-signed test. (c) Percent of syntenic (blue) and non-syntenic (orange) maize genes with a mapped eQTL, pQTL, or phosQTL (i.e., percent of genes that are a QTL trait). ** denotes *p* < 0.0001, Chi-squared test. (d) MapMan enrichment analysis of eQTL (circle), pQTL (triangle), and phosQTL (diamond) traits. Size represents number of traits. Heatmap color represents false discovery rate (FDR) from hypergeometric test for enrichment.

We wanted to explore these multi-omics data by mapping genomic regions which could regulate transcript, protein, and/or phosphosite levels. Thus, we constructed a genetic linkage map of the B73xMo17 combined genome based on genotyping data from 247 IBMDHLs consisting of 4,191 genetic markers with an average spacing of 0.4 centiMorgan (cM) for QTL mapping (Supplemental Figure 1, Supplemental Dataset 4). This map was further trimmed to account for the 134 IBMDHLs used in this study, resulting in a genomic map of 3,409 markers (Supplemental Dataset 4). Gene-products expressed in at least 50% (67 of the 134) of the lines were used as input for molecular QTL mapping. We identified 34,056 eQTL, 5,321 pQTL, and 4,935 phosQTL, with an average effect size (proportion of variance) of 3.68%, 3.50%, and 3.75%, respectively. (Figure 1a, Supplemental Dataset 5). Further, we found that *cis* QTL have a significantly higher effect size than *trans* QTL (4.50% versus 3.31%; 4.01% versus 3.33%; and 4.23% versus 3.63%, respectively) (*p* < 0.0001, Wilcoxon rank-signed test) (Figure 1b). Given that we would expect an effect size of approximately 0.029% at random (from 3,409 genetic markers), our mapped QTL contribute significantly to the genotypic variation of these gene-products.

We next examined overlap between our mapped QTL (Figure 1a). First, we observed a 25% overlap between pQTL and eQTL, which is significantly higher than what is expected by random chance (*p* < 0.0001, hypergeometric test). The significant overlap between pQTL and eQTL is present in *cis* QTL (*p <* 0.0001, hypergeometric test) but not in *trans* QTL (*p* = 0.806, hypergeometric test). Further, the only significant overlap between phosQTL and the other molecular QTL is between *cis* phosQTL and eQTL (35% overlap, *p* < 0.0001, hypergeometric test). The moderate to low degree of overlap between these different molecular QTL illustrates that while some cognate gene-products are regulated similarly, complex regulation occurs at various layers of gene expression that cannot be fully understood by examining the transcriptome alone.

Syntenically conserved maize genes are more likely to exhibit altered phenotypes when mutated (*20*). Thus, we obtained a list of syntenic orthologous genes between maize and sorghum (*21*) (Supplemental Dataset 6) and determined how many of them had at least one mapped molecular QTL (i.e., are a trait for at least one type of molecular QTL). We found that significantly more syntenic genes are molecular QTL traits than non-syntenic genes, regardless of the gene-product measured (44.9% versus 14.2%; 10.7% versus 2.9%; and 6.7% versus 1.1% for eQTL, pQTL, and phosQTL, respectively) (*p* < 0.0001, Chi-squared test) (Figure 1c). We further complemented this analysis by examining highly connected network hubs, which are more likely to be required for network integrity and organism survival than nonhubs (*22–24*). We used the STRING database (*25*) to build known protein-protein interaction networks for our molecular QTL traits (Supplemental Figure 2, Supplemental Dataset 7). We found that, for both eQTL and pQTL, the proteins with a mapped molecular QTL had a statistically significantly higher node degree (number of edges for that protein) than proteins without a mapped molecular QTL (*p* < 0.0001, Wilcoxon rank-signed test), suggesting that QTL-mediated gene-products are more likely to be protein-protein interaction network hubs. We did not find this trend for phosQTL (*p* > 0.05, Wilcoxon rank-signed test): however, this could be due to the much smaller number of phosQTL traits (198) in the protein-protein interaction network compared to eQTL (2768) and pQTL (562). Combined, these results suggest that syntenic genes and protein-protein interaction network hubs, which are both more likely to be biologically important, are genetically controlled and, further, we can map the genomic regions which control the levels of their gene-products.

We used MapMan4 (*26*) to construct an ontology for the B73-Mo17 combined genome and perform ontology enrichment analysis on our molecular QTL traits (Figure 1d, Supplemental Dataset 8) First, we found a significant overlap in enriched terms between eQTL and pQTL (34.3%, *p* < 0.0001), eQTL and phosQTL (24.8%, *p* < 0.0001), and pQTL and phosQTL (23.6%, *p* < 0.0001, hypergeometric test). However, we did not find a significant overlap between all three molecular QTL (7.5%, *p* = 1.00, hypergeometric test). Multiple terms related to plant-pathogen interactions were enriched across the different molecular QTL including pathogen response, temperature response, response to various plant hormones, autophagy, and kinase signaling.

It is well-known that much of plant immune signaling and pathogen response is realized by primary and specialized metabolites (*27–29*). Thus, we performed metabolomics and metabolic feature QTL mapping on the NLB-infected tissue collected from the 134 IBMDHLs. We detected 36,943 metabolic features (243 of which are known metabolites) (Supplemental Dataset 9) and mapped 11,393 metabolite QTL (mQTL) (158 of which were mapped for known metabolites) (Supplemental Dataset 5). We found a statistically significant overlap between our mQTL for known metabolites and each of our molecular QTL (54.4%, 56.4%, and 56.6% for eQTL, pQTL, and phosQTL, respectively; *p* < 0.0001, hypergeometric test), supporting that these altered signaling pathways result in differential metabolite levels.

### Integration of molecular and disease QTL

We next integrated our molecular QTL with disease severity data. We mapped dQTL from Area Under the Disease Progression Curve (AUDPC) measurements (*30*) taken over six weeks from 247 IBMDHLs grown in Aurora, NY (years 2011 & 2012) and Boone, IA (years 2014 & 2015). (Figure 2A, Supplemental Dataset 10). We integrated our mapped dQTL with previously published NLB dQTL (*31*) to form a comprehensive set of 49 NLB dQTL (Figure 2B, Supplemental Table 1). We found that a significantly higher number of dQTL were mapped to chromosomes 1, 2, 4, and 8 than expected by random chance (n = 9, 11, 6, and 6 dQTL, respectively; *p* < 0.0001, hypergeometric test). Further, we found that 54.4%, 56.4%, 56.6%, and 50.0% of eQTL, pQTL, phosQTL, and mQTL, respectively, overlap with these dQTL and represent a significant enrichment (*p* < 0.0001, hypergeometric test) (Supplemental Dataset 5). This degree of overlap between these molecular and dQTL suggests our dataset will be useful for uncovering genetic determinants of NLB resistance. Furthermore, this result also highlights that these data will be useful for understanding non-stress regulatory events related to maize leaf growth and physiology.

**Figure 2.**
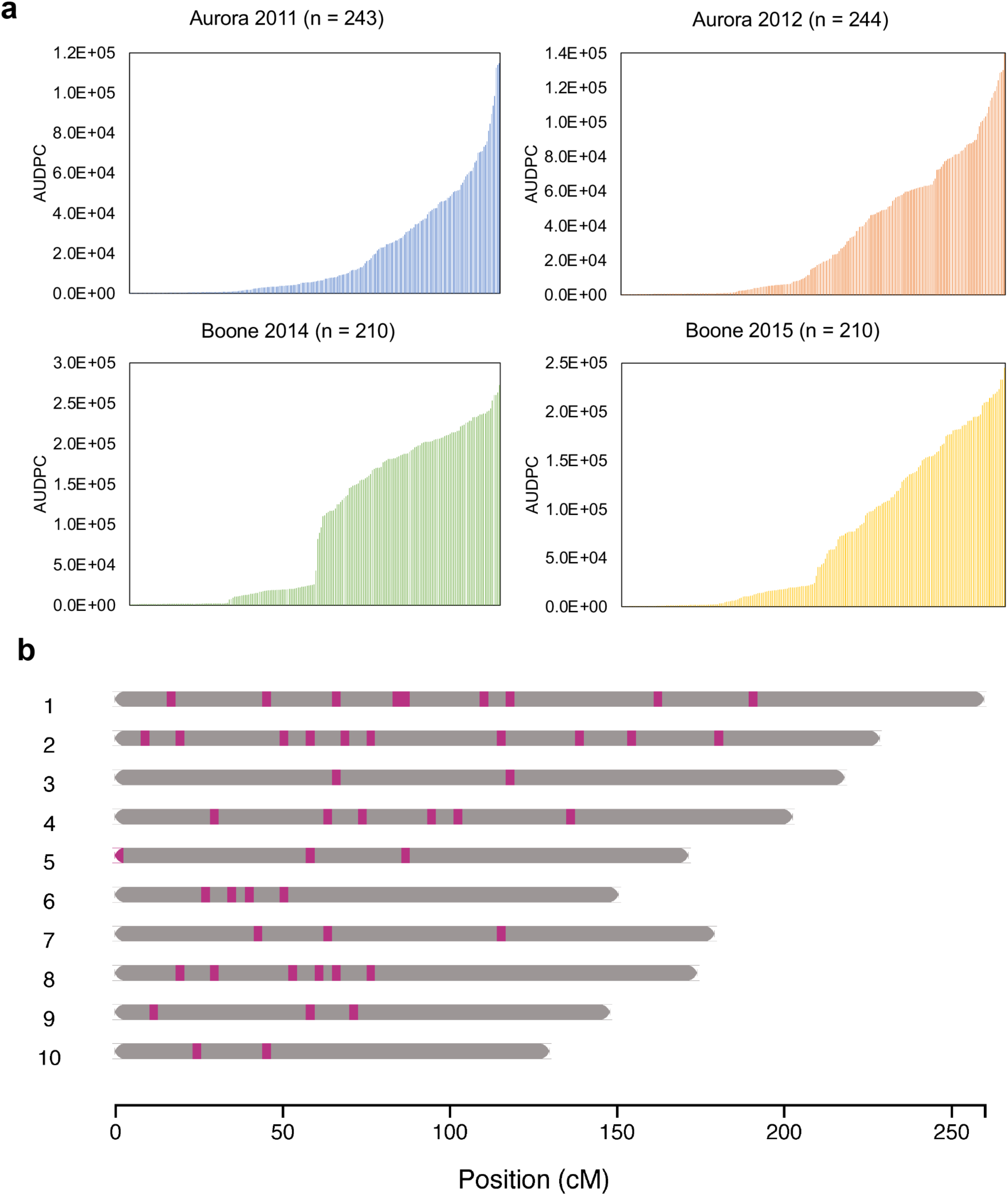
Disease QTL mapping of the NLB-infected IBMDHL population. (A) Distribution of Area Under the Disease Progression Curve (AUDPC) scores for IBMDHLs planted in Aurora 2011 & 2012 (top row) and Boone 2014 & 2015 (bottom row). Each column represents one IBMDHL. (B) Chromosome map of disease QTL for NLB response. Red lines represent the midpoint of each disease QTL.

We additionally performed transcriptomic and (phospho)proteomic profiling on four biological replicates of the B73 and Mo17 parental lines that were or were not infected with NLB (Supplemental Dataset 11). We found that there is a significant overlap between molecular QTL traits and gene products differentially expressed (DE) in response to infection in the parental lines (26.8%, 38.7%, and 40.1% overlap for eQTL, pQTL, and phosQTL, respectively) (*p* < 0.0001, hypergeometric test). This suggests that a significant proportion of our molecular QTL traits exhibit disease responsive gene expression changes.

### Integrative omics networks predict the molecular signaling cascade in response to NLB

While our molecular and dQTL mapping provide information about the genomic regions which mediate molecular NLB response and disease phenotypes, we are still missing information about the downstream signaling events, such as kinase-signaling, transcription factor (TF)-target regulation, and metabolite production. Thus, we incorporated systems genetics analyses to build regulatory networks for our mapped QTL as well as their regulated traits. We specifically leveraged our integrative omics network inference method Spatiotemporal Clustering and Inference of Omics Networks (SC-ION) (*32*) to predict a comprehensive molecular network. This integrative omics approach was used because it has been shown that networks inferred using different gene-products have unique topologies, and therefore different conclusions regarding the signaling events regulating a biological process may be observed (*32–34*). Further, integrating networks inferred from multiple gene-products improves the overall network prediction and improves the understanding of information flow in a given system (*33*).

We first used SC-ION to predict TF-centered networks for all of the mapped eQTL and pQTL (Supplemental Dataset 12). The first step of SC-ION clusters gene-products into subgroups and then infers one network for each cluster. Typically, SC-ION uses an unsupervised approach, such as clustering gene-products based on their expression in different tissues or time points. In this instance, we leveraged our molecular QTL to create supervised clusters for our network inference approach, inferring one network per eQTL or pQTL. This approach is useful because it leverages the molecular QTL mapping in the inferred networks and predicts the causal TFs in the QTL intervals, which can help identify key regulators underlying large QTL intervals that may contain tens to hundreds of putative regulators. Additionally, we leveraged the integrative omics framework of SC-ION by inferring two complementary networks: one where protein abundance (or transcript abundance when the cognate protein was not detected) is used for TF expression, and another where phosphosite intensity is used (Figure 3A). The choice of abundance or phosphosite data resulted in networks with different regulatory topologies, which is supported by other studies (*32–34*). Finally, since pQTL traits could be regulated by biological processes other than transcription, we inferred two additional pQTL networks where all of the proteins in the QTL interval could be potential regulators, not just transcription factors.

**Figure 3.**
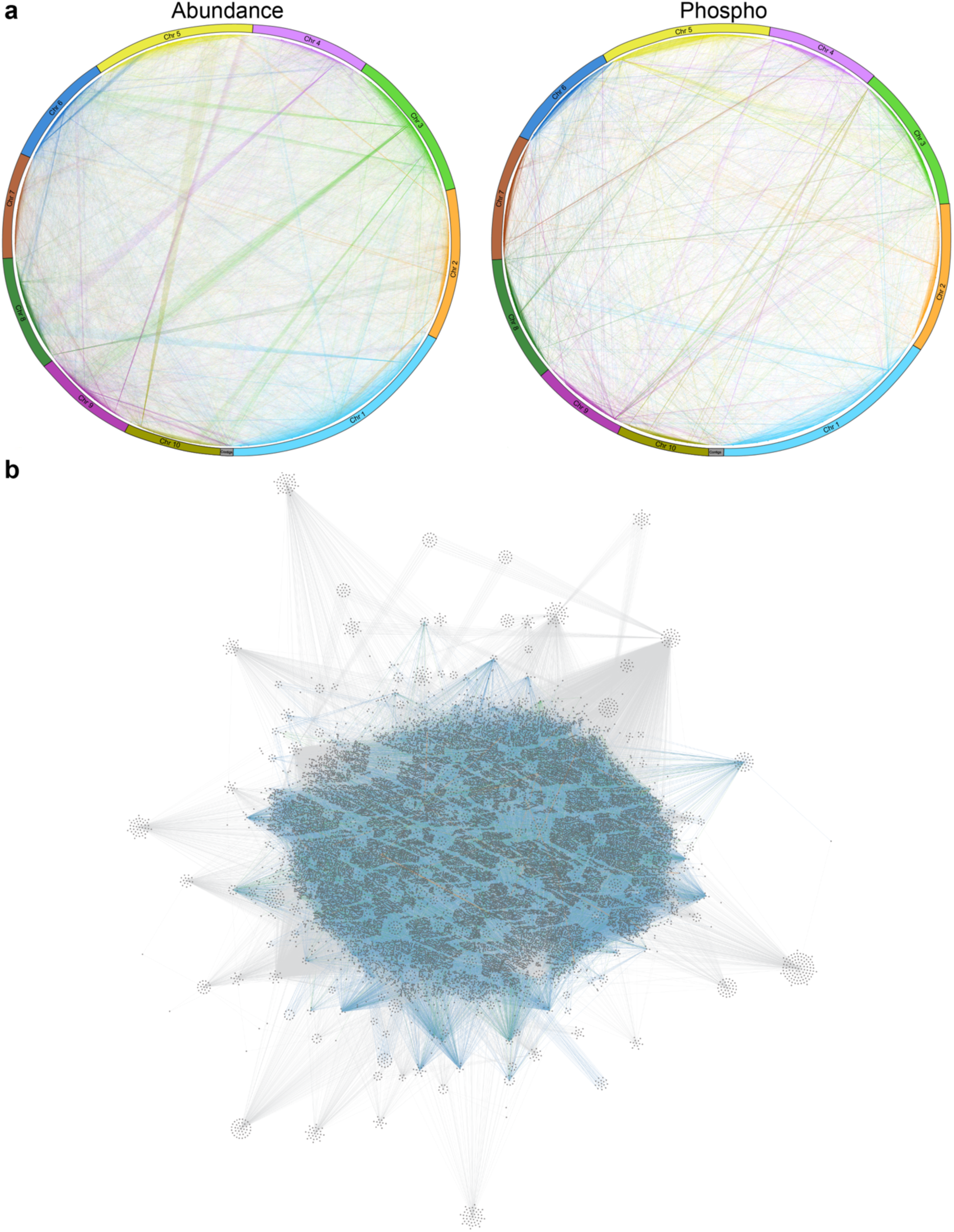
Integrative omics networks of NLB response. (a) SC-ION generated networks for eQTL using abundance (left) or phosphosite (right) data. Genes are arranged in a circos plot, beginning on the bottom right (chr1, light blue) and continuing counterclockwise. Edge color represents chromosomal location of regulator. Gray regions are contigs. (b) SC-ION generated integrated omics network for all molecular QTL. Dark gray circles (nodes) are genes. Edge color represents QTL category: eQTL (blue), pQTL (green), phosQTL (orange), mQTL (gray). The network is arranged in an organic layout, where the top of the network is towards the center, and the bottom of the network is towards the edges.

To predict kinase-signaling networks, we leveraged a correlation-based approach using kinases with an annotated activation loop, or p-loop domain (*35*, *32*, *36*). We correlated the phosphosite intensity of these p-loop domains detected in this experiment with all detected phosphosites. We then incorporated phosQTL predictions by keeping only the edges that were also predicted by our phosQTL mapping (Supplemental Dataset 12). Finally, we used SC-ION to predict the causal genes for each of the 237 known mQTL as we did for the eQTL and pQTL, although in this case we did not restrict the list of regulators to only TFs. This formed a comprehensive integrative omics network, beginning from kinase-signaling and ending with differential metabolite regulation (Figure 3B, Supplemental Dataset 12).

### Identification of candidate kinases from a phosQTL hotspot

Given the importance of kinase signaling and phosphorylation events in known plant immune signaling networks (*37*, *38*) we leveraged our phosQTL mapping to identify putative kinases responsible for alterations in phosphorylation events upon infection with NLB. We identified 56 phosQTL hotspots in our dataset with the majority (24 hotspots, 42.8%) occurring on chromosome 1 (Figure 4, Supplemental Table 2). We further identified three hotspots, two on chromosome 1 and one on chromosome 6, that contained less than three annotated kinases. Our rationale behind focusing on these three hotspots is that one of these few kinases maybe the causal kinase for differential phosphorylation events in that QTL interval. We selected one kinase from each of these intervals (Zm00001d027934, LTK1; Zm00001d029964; and NSK1) and determined their potential targets using Multiplexed Assay for Kinase Specificity (MAKS) (*39*, *36*) (Supplemental Figure 3).

**Figure 4.**
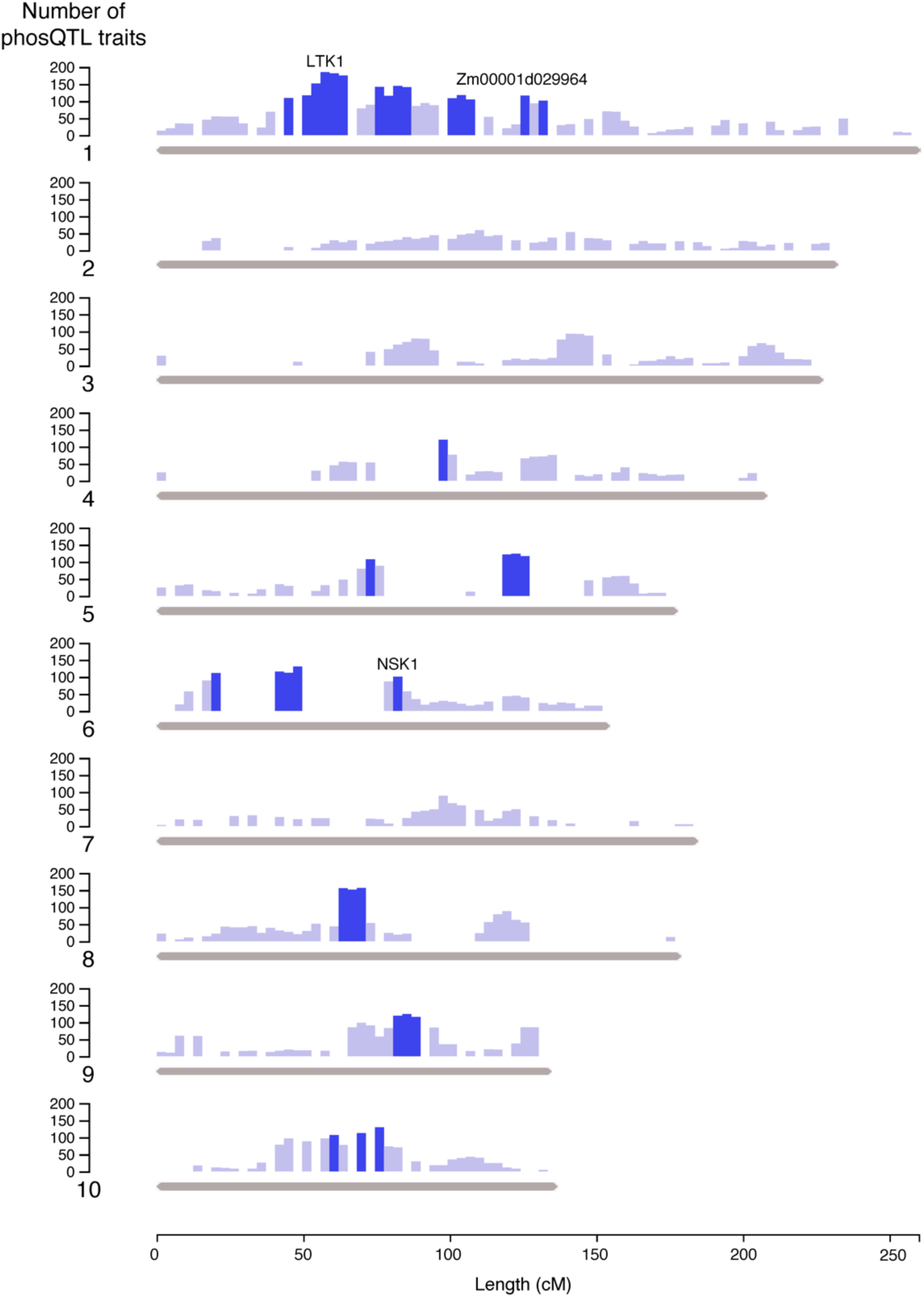
Genomic positions of phosQTL hotspots. Bar height represents number of phosQTL traits mapped to each genetic interval. Dark blue bars represent phosQTL hotspots. Labels mark the positions of phosQTL hotspots containing LTK1, Zm00001d029964, and NSK1.

Using MAKS, we identified 400, 891, and 634 substrate proteins (536, 1251, and 839 phosphosites) for LTK1, Zm00001d029964, and NSK1, respectively. Further, 34.5% to 37.5% of the kinase-specific phosQTL traits overlapped with these MAKS- enriched substrates, representing a statistically significant overlap (LTK1: *p* < 0.01; Zm00001d029964: *p* < 0.10; NSK1: *p* < 0.05, hypergeometric test) (Supplemental Dataset 13). These results show that predicted phosQTL traits are enriched in the MAKS targets, supporting that these kinases are responsible for phosphorylation of the phosQTL trait proteins. We further leveraged our MAKS results to perform candidate phosQTL mapping, which mapped an additional 59, 104, and 128 phosQTL for LTK1, Zm00001d029964, and NSK1, respectively. In total, we identified 19, 20, and 57 MAKS- enriched phosphoproteins (36, 44, and 102 MAKS-enriched phosphosites) regulated by the hotspot phosQTL containing LTK1, Zm00001d029964, and NSK1, respectively (Supplemental Dataset 13). We found that 77 (80.2%) of these phosphoproteins are uniquely enriched for one kinase (and therefore one phosQTL), suggesting that these three kinases control different signaling pathways in response to NLB.

### NSK1 and bHLH106 modulate NLB response through brown midrib2

Based on our MAKS results, we became particularly interested in the kinase NSK1. NSK1 had the largest number of MAKS-enriched direct targets which overlapped with mapped phosQTL (57) of the three kinases we tested. Further, NSK1 was one of the mapped genes with an associated SNP from the previously published large-scale NLB dQTL study (*31*). We thus used our integrative omics network to predict the downstream signaling events of NSK1 and its MAKS targets (Supplemental Figure 4).

We annotated the network using a curated list of 255 genes implicated in NLB resistance which were identified through GWAS, recombinant inbreed line (RIL) mapping, and/or mutant analysis (*31*, *40–46*, *2*) (Supplemental Table 3) and examined if any of these genes were predicted to be downstream of NSK1. We found that one of NSK1’s targets, a TF annotated as *bHLH106* (Zm00001d020834), is predicted to regulate *BM2* (Zm00001d034602) (Supplemental Figure 4). Furthermore, *bm2* mutants display heightened susceptibility to NLB, which wasn not detected via association analysis (*2*). This led us to further investigate if a pathway between NSK1, bHLH106, and BM2 could modulate NLB resistance.

To validate the predicted signaling pathway between NSK1, bHLH106 and BM2, we performed a transient protoplast assay where a construct with the *BM2* promoter driving luciferase (*Pro_BM2_:LUC*) was co-expressed with a construct constitutively expressing *NSK1* (*Pro_CsVMV_:NSK1*) or *bHLH106* (*Pro_CsVMV_:bHLH106*). Both NSK1 and bHLH106 co-expression resulted in higher luciferase activity compared to the *BM2* promoter alone (Figure 5A), suggesting that *BM2* is a downstream target of both NSK1 and bHLH106. To further investigate the mechanism of NSK1 and bHLH106 in NLB resistance, we generated CRISPR-induced mutant lines of each (Supplemental Figure 5, see Methods) and measured the expression of *BM2* in the mutants and corresponding WT controls with NLB infection or mock treatment. Our results show that, when compared to WT, *nsk1* and *bhlh106* mutants have significantly lower levels of *BM2* transcript only after NLB infection (p<0.05, two-sample t-test) and not with mock treatment (Figure 5B-C and Figure 5F-G). Finally, we performed NLB disease infection assays on *nsk1* and *bhlh106* mutants by inoculating 4-week old plants with NLB and measuring the lesion area on each plant after 13 days. We found that both *nsk1* and *bhlh106* mutation are more susceptible to NLB compared to a WT control based on larger legion areas after 13 days (p<0.05, two-sample t-test) (Figure 5D-E and Figure 5H-I). These results validate our network-inferred prediction of a novel signaling pathway incorporating a kinase (NSK1), a transcription factor (bHLH106), and a lignin biosynthesis enzyme (BM2) to promote NLB resistance (Figure 5J).

**Figure 5.**
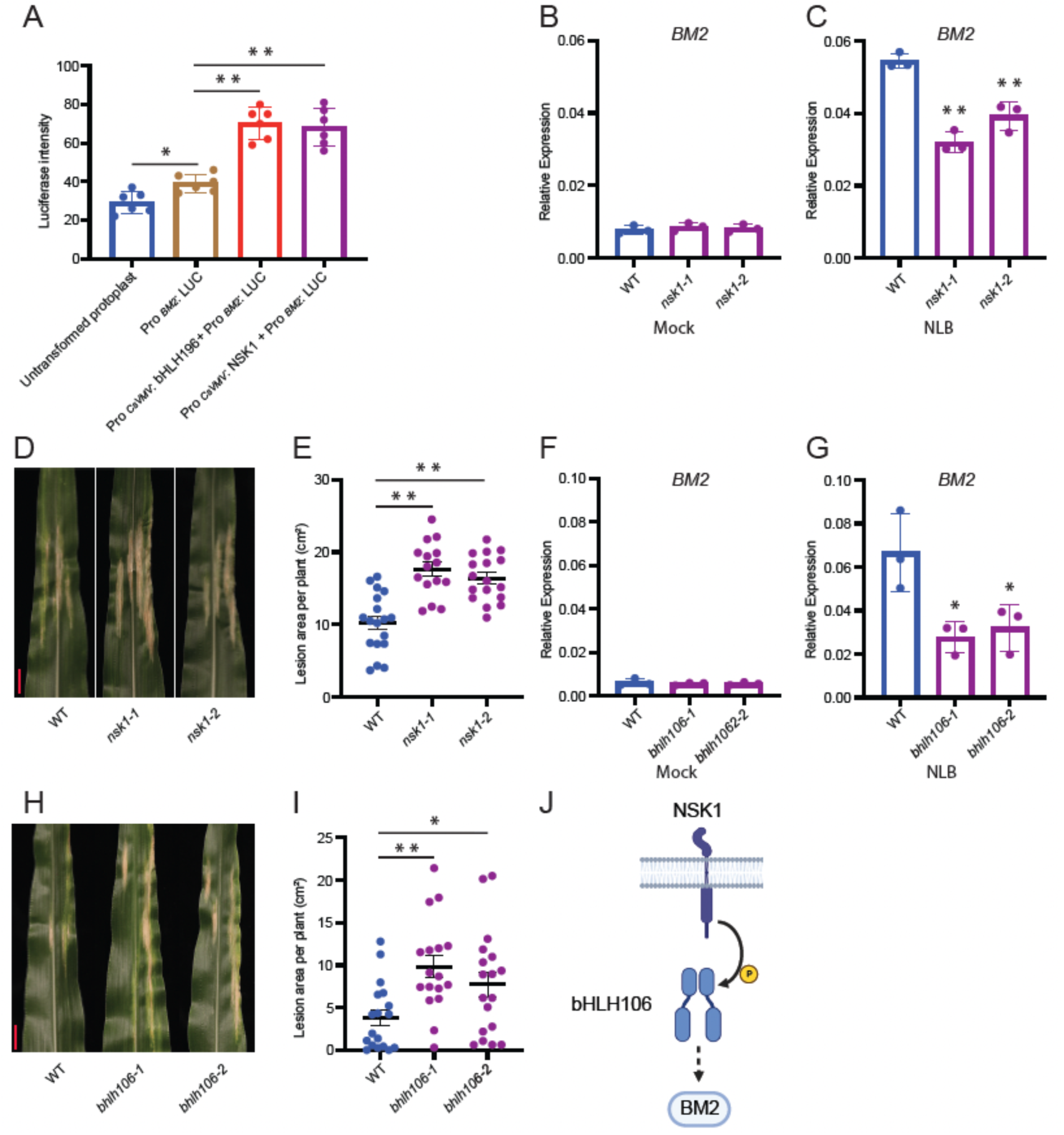
NSK1 and bHLH106 are required for resistance to NLB. (A) Maize protoplast transient assay testing *BM2* promoter activity when co-expressed with NSK1 or bHLH106 (n=6). RT-qPCR measuring *BM2* expression in mock (B) or NLB infected (C) *nsk1* and WT plants (n=3 for each sample). (D) Leaf images of the lesion region 13 days post inoculation for *nsk1-1*, *nsk1-2,* and WT. (E) Data (lesion area per plant) analysis of NLB infection assay with *nsk1-1* (n=15), *nsk1-2* (n=17), and WT (n=18). (F) RT-qPCR measuring *BM2* expression in mock (F) or NLB infected (G) *bhlh106* and WT plants (n=3 for each sample). (H) Leaf images of the lesions region 13 days post inoculation for *bhlh106-1*, *bhlh106-2,* and WT. (I) Data (lesion area per plant) analysis of NLB infection assay with *bhlh106-1* (n=17), *bhlh106-2* (n=18), and WT (n=18). *denotes *p* < 0.05, \*\**p* < 0.01, t-test. Scale bar = 2 cm. (J) Model for NLB resistance mediated by NSK1 and bHLH106 induction of *BM2*.

### Conclusions

Here, we produced a unique dataset incorporating five different QTL sources (eQTL, pQTL, phosQTL, mQTL, and dQTL) to identify novel signaling pathways regulating NLB response in corn. Importantly, the incorporation of a large-scale phosQTL dataset allowed us to more deeply investigate the role of phosphorylation in plant immune response. Our combination of MAKS with integrative omics network inference specifically identified a regulatory pathway connecting the kinase NSK1, the TF *bHLH106*, and lignin biosynthesis enzyme BM2. Additionally, our large-scale datasets and integrative omics networks provide a resource for the greater plant immune signaling community to identify and investigate putative regulatory mechanisms. Although our networks and data are NLB-specific, the high degree of overlapping dQTL between different plant diseases (*47*) suggests that some of these pathways may regulate disease response more generally. In conclusion, our work demonstrates the importance of incorporating the effect of post-translational modifications, such as phosphorylation, with quantitative genetics approaches to identify novel pathways controlling biological processes.

## Acknowledgements

We dedicate this manuscript to Dr. Nick Lauter (1972-2021), whose exceptional contributions were instrumental to this study. We thank Miriam Lopez (USDA ARS) for all her contributions in field planting and scoring disease severity for QTL mapping. This work was supported by funds from USDA Hatch Act, ISU Plant Sciences Institute, and NSF (IOS-1759023) to JWW and a USDA-NIFA Postdoctoral Research Fellowship (2019-67012-29712) to NMC.

## Data Availability

Mass spectrometry data can be downloaded from MassIVE (http://massive.ucsd.edu) using the identifiers MSV000094728, MSV000094729, and MSV000094768. Transcriptome data is available at NCBI Sequence Read Archive (SRA) using the BioProject accession number PRJNA1129186.

## Methods

### Tissue collection for multi-omics profiling

The Intermated B73xMo17 Doubled Haploid Lines (IBMDHLs) used in this study were acquired from Pioneer-HiBred. They were developed by crossing the B73 and Mo17 parental lines, followed by random mating for 10 subsequent generations and double haploidization (*1*). Parental B73 and Mo17 inbreds as well as one hundred and thirty-four IBMDHLs and the B73 and Mo17 parental lines were planted at the Agronomy Research Farm in Boone, Iowa, USA in 2018 in a randomized complete block design. Four-week-old plants (V6-V7) were whorl- inoculated with approximately eight sorghum grains colonized with the NLB-IA01 isolate and 1 ml water was then added to each whorl. Tissue from the NLB infected (the region around infection flecks) primary leaf was collected seven days post-inoculation. Infected leaf tissue from three different plants was pooled for each IBMDHL and used for transcriptomic, proteomic, phosphoproteomic, and metabolomic analysis.

### Transcriptomic analysis

Total RNA was extracted from 100 mg tissue for each sample using Trizol (Invitrogen) and RNeasy mini kit (Qiagen). 500 ng of RNA was used with the QuantSeq 3’ mRNA-Seq Library Prep FWD Kit for Illumina (Lexogen) to construct libraries. Sequencing was performed on a HiSeq 3000 with 100bp single-end reads. Libraries were run twice to increase the number of total reads to ∼6 million per sample. The combined B73-Mo17 genome was constructed by using gmap (*2*) to merge the B73 v4 (*3*) and Mo17 CAU (*4*) genomes. Adapters were trimmed using ea-utils (*5*), reads were mapped to the combined B73-Mo17 genome using the STAR aligner (*6*), alignments for the two runs were merged using samtools (*7*, *8*), and read counts were obtained using htseq-count (*9*). Read counts were normalized using Trimmed Means of M (TMM) normalization (*10*). The TMM-normalized gene expression values were used as input for QTL mapping (Supplemental Dataset 1).

### Protein extraction

Protein extraction and digestion were done using Phenol-FASP (*11*, *12*). Briefly, the leaf tissue was ground under liquid nitrogen for 15 minutes and saved at -80°C until use. 150 mg ground tissue for each sample was used for protein extraction and mixed with 5 volumes of Tris buffered phenol pH 8 (buffer: tissue, v: w) and 5 volumes (buffer: tissue, v: w) of sucrose buffer (50 mM Tris pH 7.5, 1 mM EDTA pH 8, 0.9 M sucrose, 1x phosphatase inhibitors(2.5 mM NAF, 250 µM NaVO4, 250 µM NaPyroPO4, 250 µM Glycerol-P), 1x HDAC inhibitors (10mM Sodium butyrate)). Proteins in phenol phase were recovered and transferred to a new tube. A second round of phenol extraction was performed on the remaining sucrose buffer. Collected proteins in the phenol phase were precipitated with 0.1 M ammonium acetate in methanol by centrifugation at 4,500 x g for 10 min at 4°C. Next, the protein pellet was probe sonicated to resuspend and re-precipitated with 0.1 M ammonium acetate in methanol, this step was repeated once more time, followed by one more resuspension and precipitation with 70% methanol. Finally, the protein pellet was placed in a vacuum concentrator till near dry and resuspended with urea resuspension buffer (8M urea, 50 mM Tris pH 7, 5 mM TCEP, 1x phosphatase inhibitors, 1 x HDAC inhibitors), the protein concentration was then determined using the Bradford assay (Thermo Scientific).

### Filter Aided Sample Preparation (FASP) and desalting

FASP was done for each sample following the methods described previously (*11*). For this, we used Amicon Ultracel – 30K centrifugal filters (Cat # UFC803008). Resuspended proteins were cleaned on filter though sequential washes with urea solution (8 M urea, 100 mM Tris pH 8, 1x phosphatase inhibitors, 1x HDAC inhibitor). Proteins were reduced with 2 mM TCEP and alkylated with 50 mM iodoacetamide during the process of FASP. Then on column digestion was done with trypsin (Roche 3708969001) (enzyme to protein ratio 1:100) in 0.05 M NH_4_HCO_3_ solution. After overnight incubation at 37°C, undigested protein was estimated using Bradford assays, then trypsin (1 μg/μl) was added to a ratio of 1:100 and an equal volume of Lys-C (0.1 μg/μl) was added to the sample and incubated for an additional 4 hours at 37°C. Clean, digested peptides were eluted from FASP columns by centrifugation, acidified to pH 2-3 with 100% formic acid, centrifuged at 21,000 x g for 20 min, and the supernatant was transferred to a new tube. Finally, samples were desalted using 100 mg Sep-Pak C18 cartridges (Waters). Eluted peptides were dried using a vacuum centrifuge (Thermo) and resuspended in 0.1% formic acid. Peptide amount was quantified using the Pierce BCA Protein assay kit (Thermo scientific).

### TMT Labeling

TMT11plex^TM^ label reagents (ThermoFisher, Lot #TD264169 and TA265136) were used to label the peptides for each sample according to a modified labeling method (*12*). Four hundred µg of C18 desalted peptides were resuspended in 400 µl of 0.2 M HEPES buffer pH 8.5 and then mixed with 0.53 mg TMT reagent that was resuspended in 160 µl dry acetonitrile. Each TMT11 labeling group included 10 individual lines of IBMDHLs or parental inbred lines (3 replicates for both B73 and Mo17), and one pooled sample (peptides from each of the IBMDHLs sample and parental lines were mixed together, took 400 µg for each labeling group, labeled with the TMT label-126) as a reference between different labeling groups. After 2-hour incubation at room temperature, 32 µl of 5% hydroxylamine were added to each tube and vortexed. The samples were incubated at room temperature for 15 minutes to quench the labeling reaction. Then, 538 µl out of 592 µl labeling solution of each sample was transferred to a new tube and mixed together, dried with a vacuum centrifuge (Thermo), resuspended with 5 ml 0.1% formic acid, adjusted pH to around 3 with 100% formic acid, then desalted using 500 mg Sep-Pak C18 cartridges (Waters). The elution from C18 column was dried with a vacuum centrifuge (Thermo), then moved forward for phosphopeptides enrichment or stored at -80°C. The rest 54 µl labeling solution of each sample was mixed together and stored at -80°C.

### Phosphopeptide enrichment

The TMT-labeled peptides were subjected to sequential enrichment using metal oxide affinity chromatography (SMOAC) using the manufacturers protocol (Thermo Scientific). First enrichment was performed with the High-Select TiO_2_ Phosphopeptide Enrichment Kit. The flow through from the TiO_2_ enrichment was dried with a vacuum centrifuge, then a second phosphopeptide enrichment was performed with the High-Select Fe-NTA Phosphopeptide Enrichment Kit (Thermo). TiO_2_ and Fe-NTA enriched phosphopeptides were not pooled and were analyzed independently by LC-MS/MS. Phosphopeptides were stored at -80°C until analysis.

### LC-MS/MS

An Agilent 1260 quaternary HPLC was used to deliver a flow rate of ∼600 nL min^-1^ via a splitter. All columns were packed in house using a Next Advance pressure cell, and the nanospray tips were fabricated using a fused silica capillary that was pulled to a sharp tip using a laser puller (Sutter P-2000). 12 μg of TMT-labeled peptides (non-modified proteome), or 15 μg TiO_2_ or Fe-NTA enriched peptides (phosphoproteome), were loaded onto 10 cm capillary columns packed with 5 μM Zorbax SB-C18 (Agilent), which was connected using a zero dead volume 1 μm filter (Upchurch, M548) to a 5 cm long strong cation exchange (SCX) column packed with 5 μm PolySulfoethyl (PolyLC). The SCX column was then connected to a 20 cm nanospray tip packed with 2.5 μM C18 (Waters). The 3 sections were joined and mounted on a Nanospray Flex ion source (Thermo) for on-line nested peptide elution. A new set of columns was used for every sample. Peptides were eluted from the loading column onto the SCX column using a 0 to 80% acetonitrile gradient over 60 minutes. Peptides were then fractionated from the SCX column using a series of 9 (25, 35, 40, 50, 60, 70, 80, 100, 1000 mM) and 8 (4, 15, 40, 45, 50, 70, 100, 1000 mM) salt steps (ammonium acetate) for the non-modified proteome and phosphoproteome analysis, respectively. For these analyses, buffers A (99.9% H2O, 0.1% formic acid), B (99.9% ACN, 0.1% formic acid), C (100 mM ammonium acetate, 2% formic acid), and D (1M ammonium acetate, 2% formic acid) were utilized. For each salt step, a 150-minute gradient program comprised of a 0–5 minutes increase to the specified ammonium acetate concentration, 5–10 minutes hold, 10–14 minutes at 100% buffer A, 15–120 minutes 10–35% buffer B, 121–140 minutes 45–80% buffer B, 140–144 minutes 80% buffer B, and 145–150 minutes buffer A was employed.

Eluted peptides were analyzed using a Thermo Scientific Q-Exactive Plus high-resolution quadrupole Orbitrap mass spectrometer, which was directly coupled to the HPLC. Data dependent acquisition was obtained using Xcalibur 4.0 software in positive ion mode with a spray voltage of 2.20 kV and a capillary temperature of 275 °C and an RF of 60. MS1 spectra were measured at a resolution of 70,000, an automatic gain control (AGC) of 3e6 with a maximum ion time of 100 ms and a mass range of 400-2000 m/z. Up to 15 MS2 were triggered at a resolution of 35,000 with a fixed first mass of 120 m/z for phosphoproteome and 120 m/z for proteome. An AGC of 1e5 with a maximum ion time of 50 ms, an isolation window of 1.3 m/z, and a normalized collision energy of 33. Charge exclusion was set to unassigned, 1, 5–8, and >8. MS1 that triggered MS2 scans were dynamically excluded for 45 or 25 s for phospho- and non-modified proteomes, respectively.

### Data analysis

The raw data were analyzed using MaxQuant version 1.6.3.3. Spectra were searched using the Andromeda search engine in MaxQuant against the combined B73-Mo17 FASTA file which was complemented with reverse decoy sequences and common contaminants by MaxQuant. The phosphoproteome “Phospho STY” was also set as a variable modification. The sample type was set to “Reporter Ion MS2” with “10plex TMT selected for both lysine and N-termini”. Digestion parameters were set to “specific” and “Trypsin/P;LysC”. Up to two missed cleavages were allowed. A false discovery rate, calculated in MaxQuant using a target-decoy strategy, less than 0.01 at both the peptide spectral match and protein identification level was required. The “second peptide” option to identify co-fragmented peptides was not used. The match between runs feature of MaxQuant was not utilized..

Sample loading (within-run) and internal reference (between run) (*13*) normalization were performed using the TMT-NEAT Analysis Pipeline (https://doi.org/10.5281/zenodo.5237316) (*14*). Differential expression analysis was not performed for the IBMDHLs – rather, the normalized values for protein abundance and phosphosite intensity were exported from TMT-NEAT and used for QTL mapping (Supplemental Datasets 2&3).

### Luciferase assay with maize protoplasts

The full length CDS sequence of *NSK1* (*Zm00001d037297*) and bHLH106 (*Zm00001d020834*) was synthesized by TWIST bioscience and inserted between the sequence of attL1 and attL2 of a gateway entry vector. The promoter region (2.7kb) of *BM2* was synthesized by AZENTA LIFE SCIENCE and inserted into a cloning vector between the restriction enzymes of Nco1 and Not1, then the gateway entry vector for *BM2* promoter, pENTR4-BM2Pro, was constructed by inserting the promoter fragment into the pENTR4 gateway entry vector between the restriction enzymes of Nco1 and Not1. The transient expression vectors of pCsVMV_GW- NSK1 and pCsVMV_GW-bHLH106 were constructed by gateway LR reactions between the entry vectors including the CDS sequence of *NSK1* or *bHLH106* and a gateway destination vector pCsVMV_GW. The luciferase reporter vector pOR_GW-BM2Pro was constructed by gateway LR reaction between pENTR4-BM2Pro and a gateway destination luciferase reporter plasmid- pOR_GW.

Maize leaf protoplasts were isolated from 12 days old etiolated B73 leaves followed the methods described previously (*15*). The plasmids used for protoplast transformation were prepared with GeneJET Plasmid Midiprep kit (Thermo scientific). 3.3 µg of plasmid of pCsVMV_GW-NSK1 or pCsVMV_GW-bHLH106 were co-transformed with 3.3 µg plasmid pOR_GW-BM2Pro into 50,000 protoplasts, or 3.3ug plasmid pOR_GW-BM2Pro alone was transformed with 50,000 protoplasts as a control, then the transformed protoplasts were incubated overnight with 1 mM luciferin at room temperature in dark. The next day, luminescence signal was detected by a Synergy H1 microplate reader (Biotek).

### RT-qPCR

The materials of *nsk1*-1, *nsk1-2*, and B104 (WT) or *bHLH106-1*, *bHLH106-2,* and B104 (WT) were planted in growth chamber with a long day conditions (16 hours light at 28°C, 8hours dark at 24°C), 7 days old seedlings of each genotype were treated with NLB-IA01 (100,000 spores/ml in 0.1% tween 20) and mock (0.1% tween 20) by spraying. After the spraying treatment, the plants were covered with a clear plastic dome to keep high moisture. Leaf tissue was collected 24 hours post the treatment. Total RNA was extracted with Trizol® (Invitrogen) and Direct-zol RNA MiniPrep kit (Zymo Research), cDNA was synthesized with LunaScript® RT SuperMix Kit (NEB). Quantitative PCR for *BM2* and *ZmGADPH* (reference gene) was done with the following program: 3 min at 95 °C, followed by 39 cycles of 10 s at 95 °C, 10 s at 60 °C and 30s at 72 °C with a CFX Connect Real-Time system (Bio-rad). ΔΔCt was used to analyze the gene expression data.

### CRISPR induced mutant generation

Two gRNA primers were designed for both *NSK1* and *bHLH106*, the vectors of pYPQ142- NSK1 and pYPQ142-bHLH106 were constructed by inserting the gRNA primers into a gateway entry vector pYPQ142(*16*). Destination vectors PKL2351-NSK1 and PKL2351-bHLH106 were constructed with pYPQ142-NSK1 or pYPQ142-bHLH106, pYPQ166 (*17*) and PKL2351(*18*) by multi-sites gateway reactions. Then the agrobacterium-mediated transformation was done with maize inbred line of B104 followed the methods described previously (*19*) at the ISU Crop Bioengineering Laboratory.

### NLB greenhouse infection assay

The T1 homozygous seeds of *nsk1-1*, *nsk1-2*, and B104 (WT) or T1 homozygous of *bhlh106-1*, *bhlh106-2*, and B104 (WT) were germinated and grown in growth chamber at long day conditions (16 hours light at 28°C, 8hours dark at 24°C), two weeks old seedlings of each genotype were transplanted into 1.5-gallon pots individually and moved to a greenhouse with the same conditions. Whorl inoculation was performed to four weeks old plants with 20,000 NLB- IA01 spores (100,000 spores/ ml in 0.1% tween 20, 0.2ml per plant) per plant. The lesion images were taken after 13 days of the treatment, lesion size was analyzed with ImageJ.

### Multiplexed Assay for Kinase Specificity (MAKS)

Gateway entry vectors for *Zm00001d029964* (pENTR4-Zm00001d029964), *LTK1* (pENTR4-LTK1; Zm00001d027934), and *NSK1* (pENTR4-NSK1) were constructed by cloning full length CDS for *Zm00001d029964* or CDS fragments for the C-terminal (after transmembrane domain to the end of protein) of these proteins for LTK1 and NSK1 into a gateway entry vector pENTR4. The plasmids pDEST17-Zm00001d029964, pDEST17-LTK1 and pDEST17-NSK1 were constructed by LR reactions between pENTR4-Zm00001d029964, pENTR4-LTK1 or pENTR4- NSK1 and a gateway destination vector pDEST17 for inducible high-level expression of N-terminally 6xHis-tagged proteins in bacterial cells. The recombined proteins 6xHis- Zm00001d029964, 6xHis-LTK1 and 6xHis-NSK1 were induced by IPTG in an *E. coli* strain BL21 and purified using nickel nitrilotriacetic acid (Ni-NTA) agarose resin. Plant protein extraction for MAKS was done using ground leaf tissue from an equal mix of B73 and Mo17 seedling tissue and the phenol-FASP protocol as described previously (*11*). Four replicates of 6xHis- Zm00001d027934, 6xHis-LTK1 6xHis-NSK1, or 6xHis-Gus (control) were used to perform the MAKS at a kinase:protein ratio of 1:75 following the methods described previously (*20*, *21*). Then, peptides from each replicate were TMT labeled, followed by phosphopeptide enrichment as described by (*21*). Pooled TMT labeled peptides were analyzed by LC-MS/MS, performed with the same methods described previously (*21*) (Supplementary Figure 2).

Statistical testing on detected phosphosites was performed using TMT-NEAT (https://doi.org/10.5281/zenodo.5237316) (*14*). No sample loading normalization was performed, as it is expected that samples incubated with an active kinase will have higher overall phosphosite intensities than samples incubated with the GUS negative control. Since sample loading normalization was not used, replicates with significantly higher or lower expression for each kinase were removed. This resulted in the removal of one replicate for each kinase (after outlier removal: 4 biological replicates for GUS; 3 biological replicates for each kinase). Phosphosites with a *p*-value < 0.05 and fold change > 1.2 were considered differentially enriched.

Additional candidate phosQTL were identified from MAKS candidates by performing the Wilcoxon test on *p*-values of phosQTL peaks within 10 cM of the candidate kinase as described in the “QTL mapping” section. phosQTL with a Wilcoxon *p*-value < 0.1 and a LOD score > 2.0 were considered significant candidate phosQTL.

### Metabolomic analysis

Frozen tissue samples were extracted and processed according methods described in (*22*). In brief, 100mg of tissue in 1.5-mL fast prep tubes were extracted with 750 μL of 10 mM of ammonium acetate and 750 μL of methanol (MeOH). Samples were mixed by vortex, homogenized at 6000 RPM for 30 seconds in a FastPrep^®^ FP 120 tissue homogenizer (Qbiogene), and then sonicated in an ambient water bath for 15 min. Samples were centrifuged for 10 minutes at 20,000 RCF at ambient temperature, then 1 mL of supernatant was transferred to individual 4 mL glass vials. The supernatant was dried under a nitrogen stream at 30°C, reconstituted in 200 μL of 0.1% formic acid in water, and vortexed for 30 seconds. Vial contents were transferred to a 1.5-mL snap-cap tube, placed on ice for 10 minutes, then centrifuged at 20,000 RCF for 10 minutes at ambient temperature. 150 μL of supernatant was transferred to glass LC vials for UHPLC-HRMS analysis. LCMS grade reagents (Thermo Fisher Scientific) were used throughout the extraction.

UHPLC-HRMS analysis was performed using a Q Exactive mass spectrometer coupled to a Vanquish LC System (Thermo Fisher Scientific, Waltham, MA, USA) by reverse phase gradient elution using an ACE Excel 2 C18-PFP column (2.1 mm X 100 mm, 2μm; part # EXL- 1010-1002U) at a column oven temperature of 25°C. With solvent A as 0.1% formic acid (FA) in water and solvent B as 0.1% FA in acetonitrile, UHPLC was performed at 0.35 mL/min as follows: 0 – 3 min: 100% A; 3 - 23 min: 100% A → 80% B; 23 – 26.5 min: 80% B; and 26.5 – 30 min: 100% A. The ESI source conditions were as follows: capillary voltage, +3kV; transfer capillary temperature, 325°C; ESI probe temperature, 350°C; sheath gas flow rate, 50; auxiliary gas flow rate, 10; sweep gas flow rate, 1; ion time detector scan speed, 0.1 scans/s; and automatic gain control target, 3E^6^. After chromatographic separation, samples were analyzed in full scan positive (injection volume 2 μL) and negative (injection volume 4 μL) ion modes at mass resolution 35000 scanning from *m/z* 70-1000. Spray voltage was 3.0 kV and the S-lens was set to 30%.

The raw acquisition data were processed using a similar workflow as described in (*22*). Identification was assigned to features by *m/z* (≤5 ppm) and retention time (<0.2 min), using our method-specific metabolite library produced from pure standards previously analyzed using the above-mentioned chromatographic gradient. Processed data were exported as a feature list containing the signal intensity for each feature in each sample. A small value (half the minimum value in the dataset) was used to replace zeros (no detection). The data were filtered to remove sample features with > 10% signal contribution from their corresponding features in the extraction blanks.

### Field trials and disease scoring

Field trials were conducted in two locations: Cornell University’s Musgrave Research Farm in Aurora, New York, USA and Iowa State University’s Agricultural Engineering and Agronomy Research Farm in Boone, Iowa, USA (*1*). A replicated, randomized complete block design was executed in two years at each location: 2011 and 2012 in Aurora, and 2014 and 2015 in Boone. Field trials in Aurora were conducted using 385 IBMDHLs, while trials in Boone were conducted using 330 IBMDHLs. A total of 244 IBMDHLs and 210 IBMDHLs in Aurora and Boone, respectively, were scored for disease severity and used for disease QTL mapping. The NLB inoculation procedure was the same as for the 2018 Boone trial and tissue infection for multi-omics, using NLB-colonized sorghum grains for whorl-inoculation between the V6-V7 vegetative stages. In Aurora, the NLB-NY01 isolate was used as previously described (*23*).

Disease severity was scored on a scale of 0 to 100% for necrotic (diseased) leaf area (Pataky et al, 1998) in one-week intervals. For the Aurora trials, disease scores were collected for three weeks beginning two weeks after the onset of anthesis. For the Boone trials, disease scores were collected for six weeks beginning at the V8 vegetative stage. The individual disease scores were used to calculate the Area Under the Disease Progression Curve (AUDPC) (*24–27*). The AUDPC scores for each IBMDHL in each year were used as input for disease QTL mapping (Supplemental Dataset 10).

### Genetic linkage map construction

IBMDHLs were genotyped by Pioneer-HiBred using the Illumina MaizeSNP50 BeadChip. Physical positions of single nucleotide polymorphism (SNPs) markers were determined by BLAST (basic local alignment search tool) results of SNP flanking sequences to the B73 RefGen_V4. BLAST-hits with an *e*-value less than 1e-10 were selected and their positions recorded (*28*). Markers were excluded from the analysis based on data quality (e.g. high percentage of missing data) and usefulness (e.g. monomorphic). A total of 4,191 physically positioned SNPs were considered for genetic map construction. Excluding 85 genotypes with an increased percentage of missing marker data and heterozygous calls, 247 genotypes and 4,191 SNPs were used for constructing a linkage map covering 1854.1cM using JoinMap4 (*29*) (Supplemental Dataset 4). The Haldane mapping function was used to calculate genetic distances (centiMorgans) between the markers ordered by their physical position (*30*). Average and maximal distance between markers was 0.4 cM and 17.0 cM respectively (Supplemental Figure 1).

### QTL mapping

Molecular QTL mapping was performed separately on the normalized QuantSeq (eQTL), (phospho)proteome (pQTL and phosQTL), and metabolome (mQTL) data (Supplemental Datasets 5). The genetic linkage map was filtered for only the markers which had genotype data in at least 95% of the 134 lines used for molecular profiling, resulting in 3,409 markers which were used in the molecular QTL mapping (Supplemental Dataset 4). Only gene-products detected in at least 50% of the 134 lines (67 lines) were used for molecular QTL mapping. Expression values were relative-rank normalized using the norm.rrank function in the DEMI R package (*31*) prior to performing QTL mapping. QTL mapping was performed using r/qtl2 (*32*) with the cross type (crosstype) as “doubled haploid” (dh) and an assumed genotyping error probability (error_prob) of 0.002. Genome scanning was performed using the scan1 function in r/qtl2, which uses Haley-Knott regression (*33*). Statistically significant QTL were identified using a permutation test with 10,000 permutations and a *p*-value < 0.1. LOD-1.5 intervals were calculated for the significant QTL. Effect size for QTL was estimated by calculating Best Linear Unbiased Predictors (BLUPs) using the scan1blup function in r/qtl2. phosQTL hotspots were classified as the phosQTL which had 100 or more regulated traits (top 10% of phosQTL) (Supplemental Table 2).

Additional pQTL were identified using the *p*-values from the corresponding eQTL. For each significant eQTL, the *p*-values for each pQTL peak within 10cM of the significant eQTL were obtained. Then a Wilcoxon test was used to determine if the *p*-value for the pQTL peak which overlapped with the significant eQTL was statistically lower than the *p*-values for all pQTL peaks detected in the 10cM interval. Significant pQTL had a Wilcoxon *p*-value < 0.1 and a LOD score > 2.0. This procedure was repeated in the opposite direction to determine additional eQTLs from pQTLs.

Disease QTL mapping was separately performed on the AUDPC measured per year (2011, 2012, 2014, 2015). Disease QTL were mapped using two data sets: first, using only the AUDPC for the 134 lines grown in the 2018 field season for molecular phenotyping (set 1), and second, using the AUDPC for all of the lines that were scored for disease resistance (244 lines for 2011 and 2012 Aurora; 210 lines for 2014 and 2015 Boone) (set 2). The same settings and statistical cut-offs were used for the disease QTL mapping as for the molecular QTL mapping. The significant QTL from sets 1 and 2 were merged with previously published NLB disease QTL (*34*) to form the final list of disease QTL (Supplemental Table 1).

### Protein-protein interaction network construction

Protein-protein interaction networks were constructed using STRING (*35*). All the molecular QTL traits (eQTL, pQTL, and phosQTL) were used as input. The v4 gene IDs were converted to v3 gene IDs for compatibility with the STRING database. Only interactions from experiments and databases with a confidence score > 500 (0.5) were kept in the final network (Supplemental Dataset 7). Network visualization and analysis was performed using Cytoscape 3.9.1 (*36*).

### B73-Mo17 differential ontology analysis

A functional ontology for the B73-Mo17 combined genome was constructed by using MapMan4 (*37*) (https://mapman.gabipd.org/) (Supplemental Dataset 8). Enriched terms were identified for each of the molecular QTL (eQTL, pQTL, phosQTL) using a hypergeometric test with *q-*value (FDR) < 0.05. Genes with no assigned ontology terms were eliminated from the enrichment analysis.

### B73 and Mo17 parental line differential gene-product analysis

Four biological replicates of the B73 and Mo17 parental lines were planted alongside the IBMDHLs in the 2018 field site, infected with NLB, and profiled for multi-omics expression (transcriptome and (phospho)proteome) as described for the IBMDHLs in the previous sections. Differential expression analysis was performed using PoissonSeq (*38*) on transcriptomics data and TMT-NEAT (*14*) on (phospho)proteomics data with a *p*-value cutoff of 0.05 (Supplemental Dataset 11).

### Integrative omics network inference

TF-centered networks were inferred using SC-ION version 2.1 (*14*) (https://doi.org/10.5281/zenodo.5237310). All genes detected in at least 50% (67) of the 134 IBMDHLs were used as network targets. All transcription factors (TFs) detected in at least 50% of the lines were used as network regulators. A list of putative TFs in the B73-Mo17 combined genome was generated using Grassius (*39*) and the MapMan4-generated annotation (*37*) (Supplemental Dataset 14). Molecular QTL information was used to cluster the genes for each network, where each cluster contained a unique molecular QTL interval. For eQTL, pQTL, and mQTL, two complementary networks were generated: one using the protein abundance of TFs, and another using the phosphosite intensity of TFs. For pQTL, abundance and phosphosite networks were also inferred where all of the genes, not just TFs, were regulators.

The kinase signaling network was inferred using the approach described in (*40*, *41*). Briefly, all protein sequences in the B73 and Mo17 genomes were searched for kinase domains. Activation loop (A-loop) coordinates were then obtained for all proteins with identified kinase domains. In total, A-loop coordinates were identified for 2,016 protein kinases in the combined B73-Mo17 genome. A correlation-based approach was used to build a network between the phosphosite intensity for these annotated A-loop domains (regulators) and all of the phosphosites detected in at least 50% of the lines (targets). The Spearman correlation coefficient was calculated for each regulator-target pair. Edges were kept if the Spearman correlation coefficient was greater than 0.5. Finally, the kinase signaling network was trimmed to only retain the edges that were supported by the phosQTL (e.g. the regulator and target must be in the same phosQTL). A network table integrating all of the TF-centered and kinase signaling networks is provided in Supplemental Dataset 12. Network visualization and analysis was performed using Cytoscape 3.9.1 (*36*).

---

**Supplemental Table 1.**
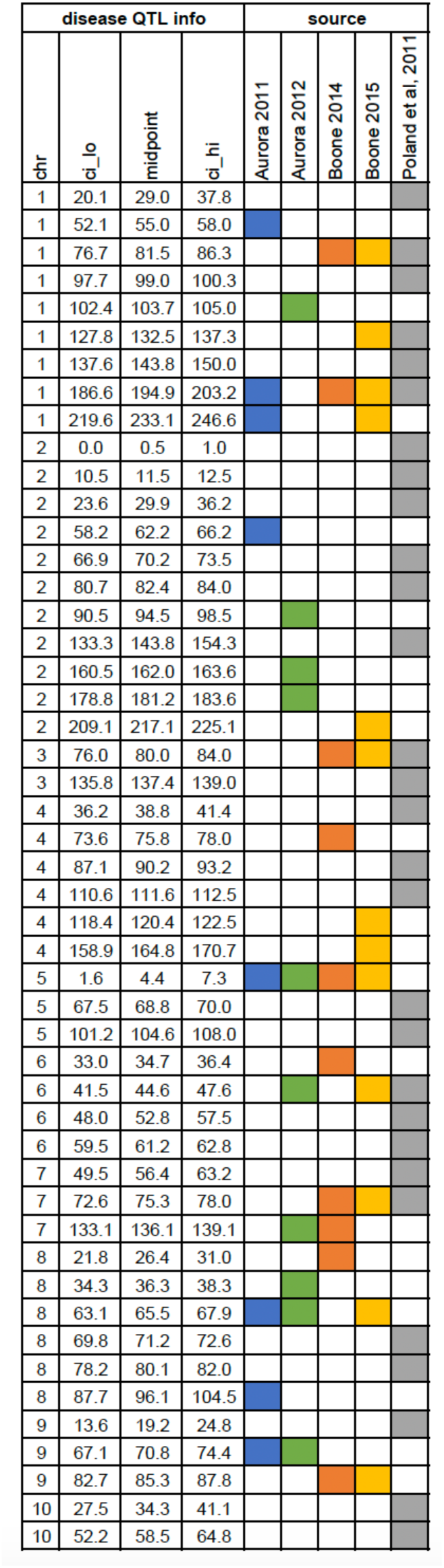
A comprehensive list of NLB disease QTL mapped in this study (Aurora 2011, blue; Aurora 2012, green; Boone 2014, orange; Boone 2015, yellow) or previously published in (*34*) (gray). chr: chromosome; ci_lo: lower 1.5-LOD interval (cM); midpoint: QTL interval midpoint (cM); ci_hi: higher 1.5-LOD interval (cM).

**Supplemental Table 2.** phosQTL hotspots identified in this study. chr: chromosome; ci_lo: lower 1.5-LOD interval (cM); ci_hi: higher 1.5-LOD interval (cM); length: length of QTL interval (cm); Number of QTL: number of phosQTL mapped to this interval.

| chr | ci.lo | ci.hi | length | Number of QTL |
| --- | --- | --- | --- | --- |
| 1 | 44.861 | 50.851 | 5.99 | 119 |
| 1 | 48.038 | 53.346 | 5.308 | 104 |
| 1 | 50.071 | 55.864 | 5.793 | 111 |
| 1 | 50.851 | 58.018 | 7.167 | 135 |
| 1 | 53.756 | 61.124 | 7.368 | 151 |
| 1 | 55.32 | 64.102 | 8.782 | 185 |
| 1 | 58.018 | 66.104 | 8.086 | 187 |
| 1 | 59.708 | 67.83 | 8.122 | 175 |
| 1 | 60.126 | 68.79 | 8.664 | 176 |
| 1 | 61.124 | 69.996 | 8.872 | 174 |
| 1 | 73.595 | 81.509 | 7.914 | 166 |
| 1 | 73.831 | 83.17 | 9.339 | 169 |
| 1 | 76.7 | 83.897 | 7.197 | 115 |
| 1 | 79.799 | 88.487 | 8.688 | 144 |
| 1 | 82.108 | 89.456 | 7.348 | 141 |
| 1 | 96.616 | 104.979 | 8.363 | 118 |
| 1 | 98.247 | 106.286 | 8.039 | 119 |
| 1 | 99.419 | 107.362 | 7.943 | 121 |
| 1 | 100.159 | 107.853 | 7.694 | 114 |
| 1 | 103.863 | 109.134 | 5.271 | 105 |
| 1 | 104.288 | 112.639 | 8.351 | 112 |
| 1 | 120.744 | 130.636 | 9.892 | 116 |
| 1 | 127.66 | 136.972 | 9.312 | 106 |
| 1 | 128.792 | 137.147 | 8.355 | 104 |
| 4 | 95.803 | 101.24 | 5.437 | 120 |
| 5 | 69.092 | 74.207 | 5.115 | 101 |
| 5 | 69.679 | 76.237 | 6.558 | 113 |
| 5 | 115.186 | 124.689 | 9.503 | 121 |
| 5 | 117.432 | 126.584 | 9.152 | 123 |
| 5 | 119.153 | 128.814 | 9.661 | 123 |
| 5 | 121.499 | 131.431 | 9.932 | 116 |
| 6 | 17.637 | 26.685 | 9.048 | 114 |
| 6 | 20.131 | 27.775 | 7.644 | 111 |
| 6 | 41.345 | 48.648 | 7.303 | 125 |
| 6 | 41.594 | 51.401 | 9.807 | 106 |
| 6 | 43.366 | 53.17 | 9.804 | 112 |
| 6 | 46.754 | 55.623 | 8.869 | 130 |
| 6 | 78.026 | 84.864 | 6.838 | 101 |
| 8 | 61.105 | 70.026 | 8.921 | 155 |
| 8 | 64.853 | 71.437 | 6.584 | 152 |
| 8 | 65.336 | 72.118 | 6.782 | 150 |
| 8 | 66.843 | 75.347 | 8.504 | 156 |
| 9 | 67.107 | 76.451 | 9.344 | 104 |
| 9 | 79.781 | 89.232 | 9.451 | 119 |
| 9 | 81.491 | 90.785 | 9.294 | 123 |
| 9 | 82.729 | 92.401 | 9.672 | 127 |
| 9 | 83.964 | 93.931 | 9.967 | 121 |
| 9 | 86.094 | 94.427 | 8.333 | 115 |
| 10 | 43.508 | 52.076 | 8.568 | 103 |
| 10 | 43.935 | 52.854 | 8.919 | 103 |
| 10 | 56.356 | 64.084 | 7.728 | 111 |
| 10 | 57.906 | 65.739 | 7.833 | 107 |
| 10 | 58.864 | 68.171 | 9.307 | 106 |
| 10 | 66.093 | 75.975 | 9.882 | 112 |
| 10 | 72.15 | 80.405 | 8.255 | 128 |
| 10 | 74.12 | 84.006 | 9.886 | 131 |

**Supplemental Table 3.**
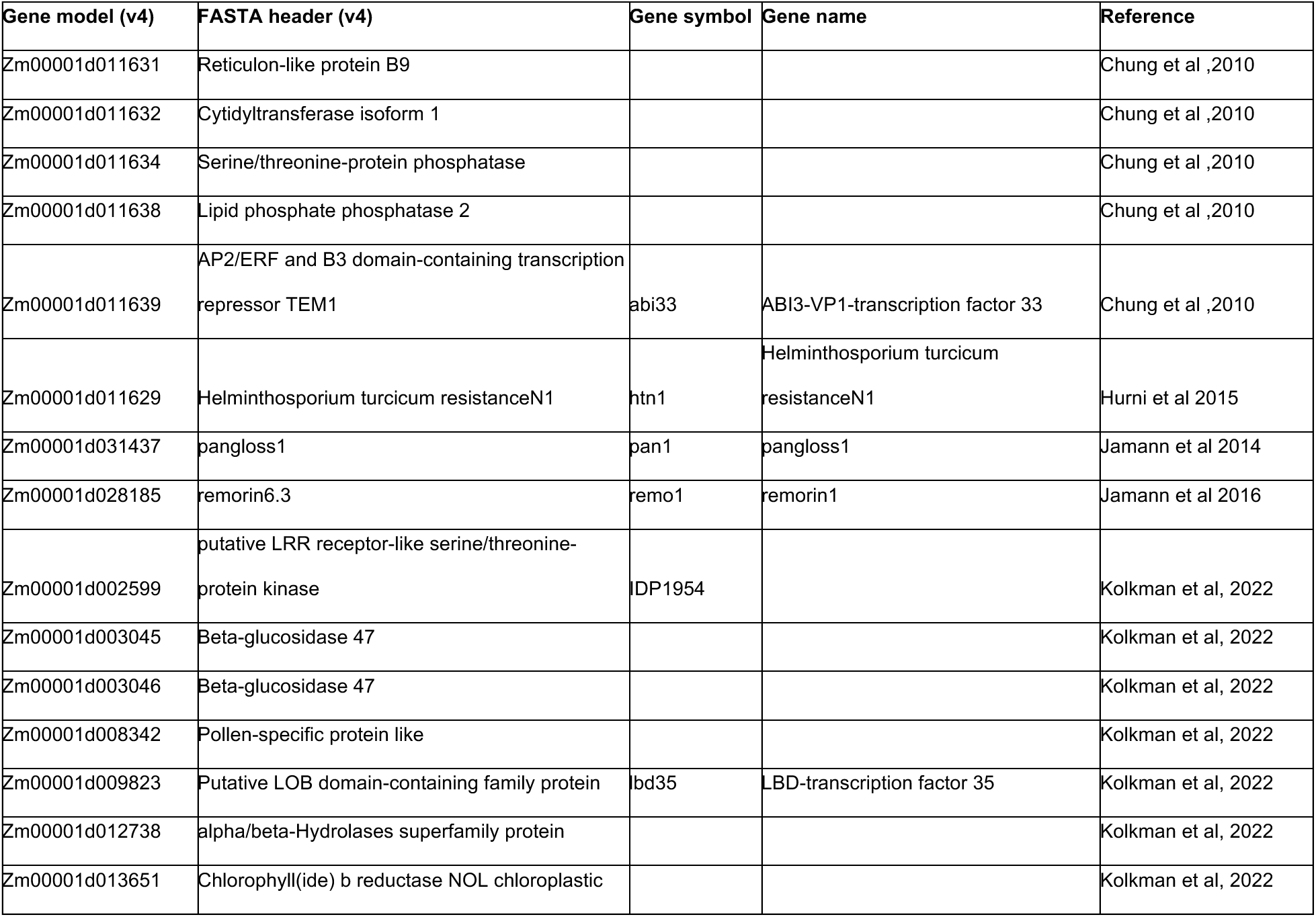
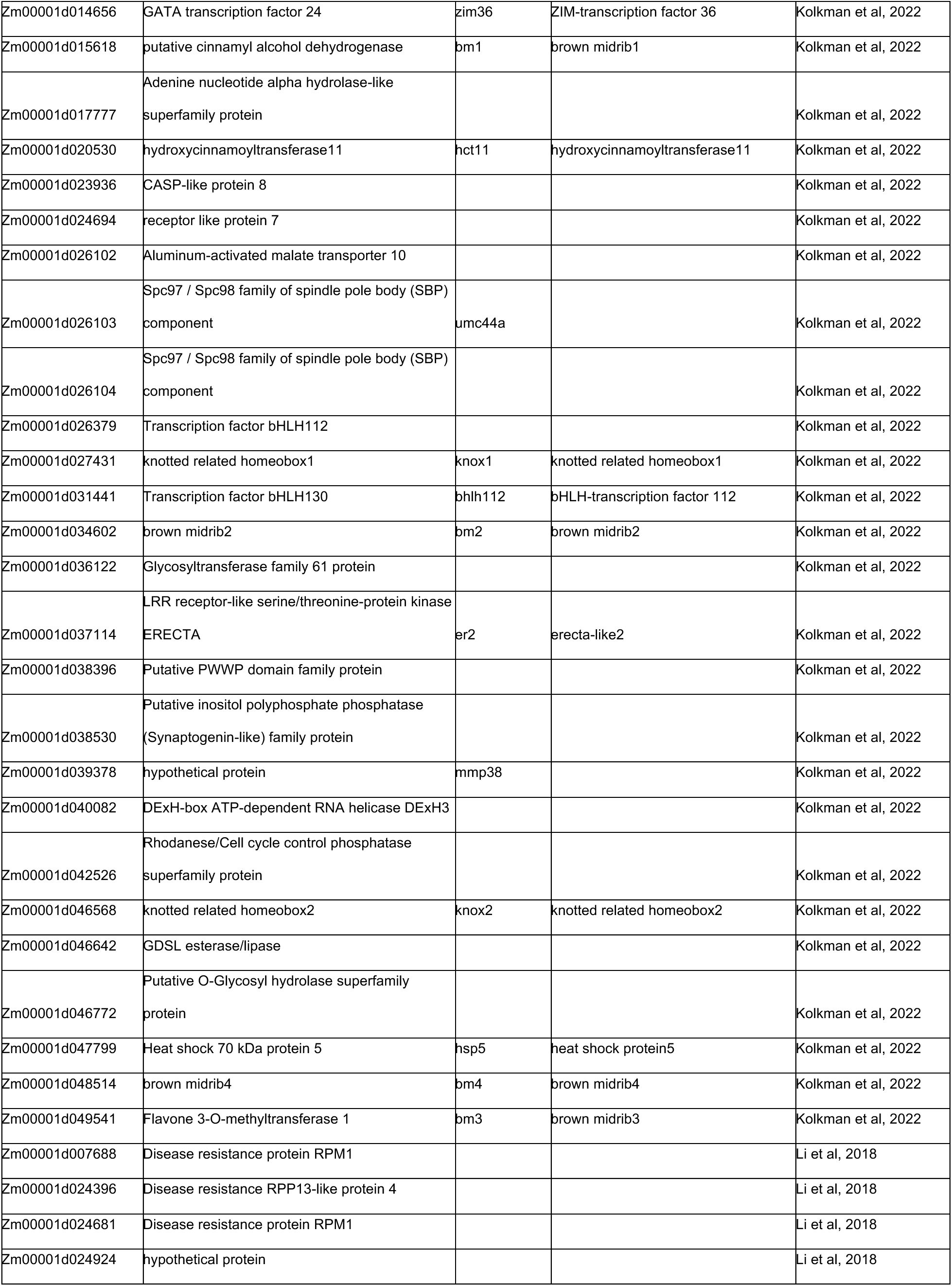
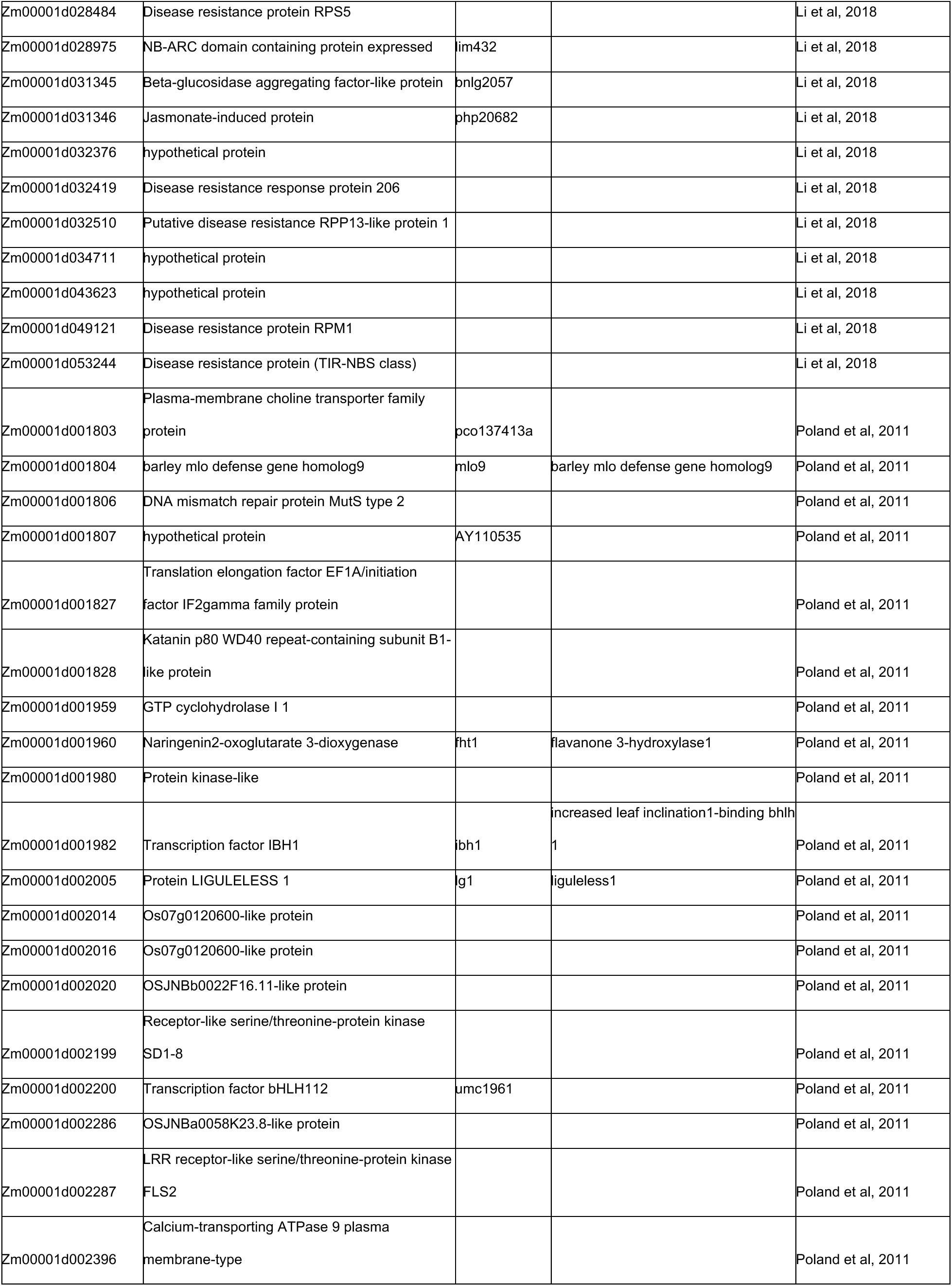
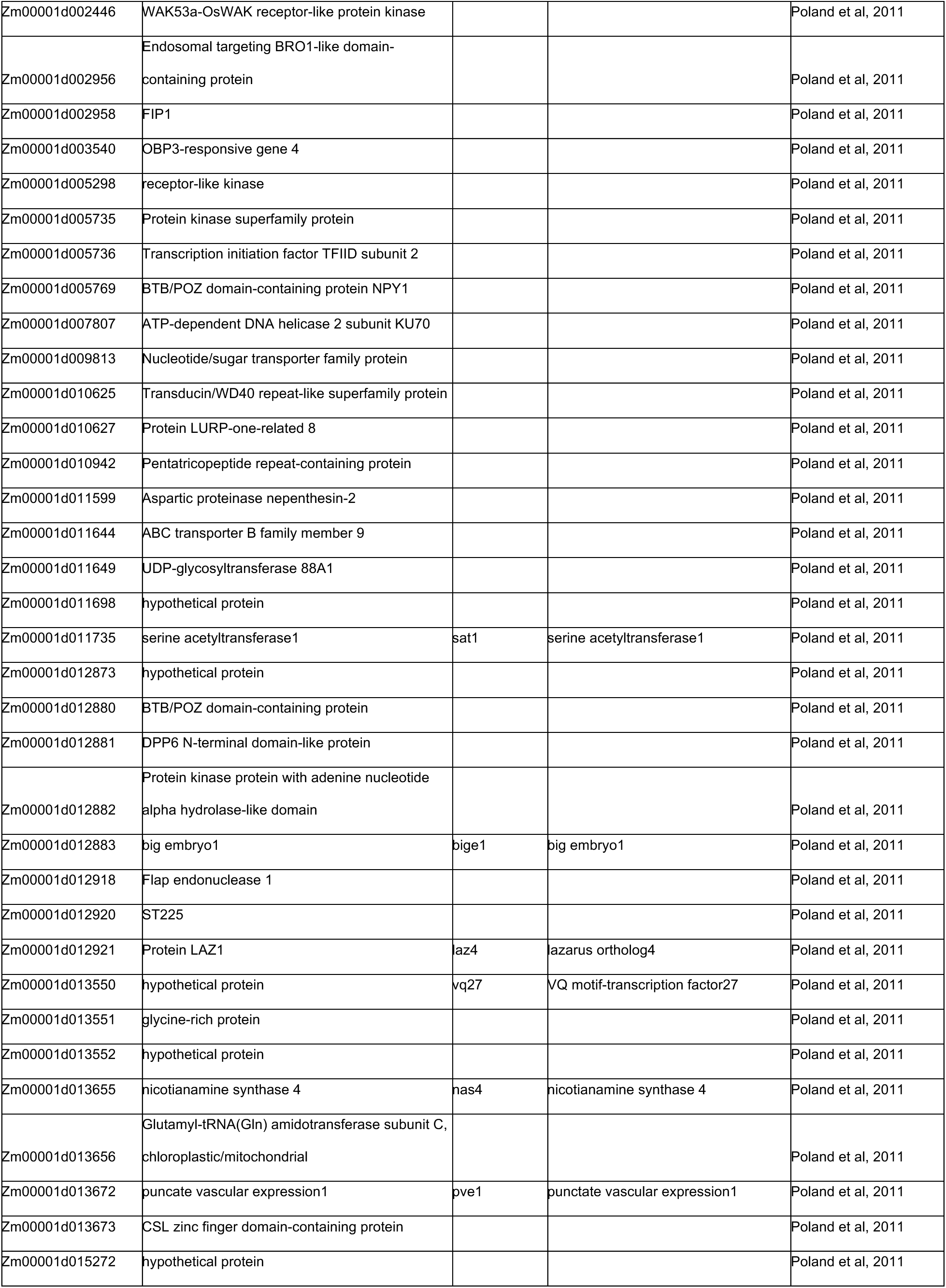
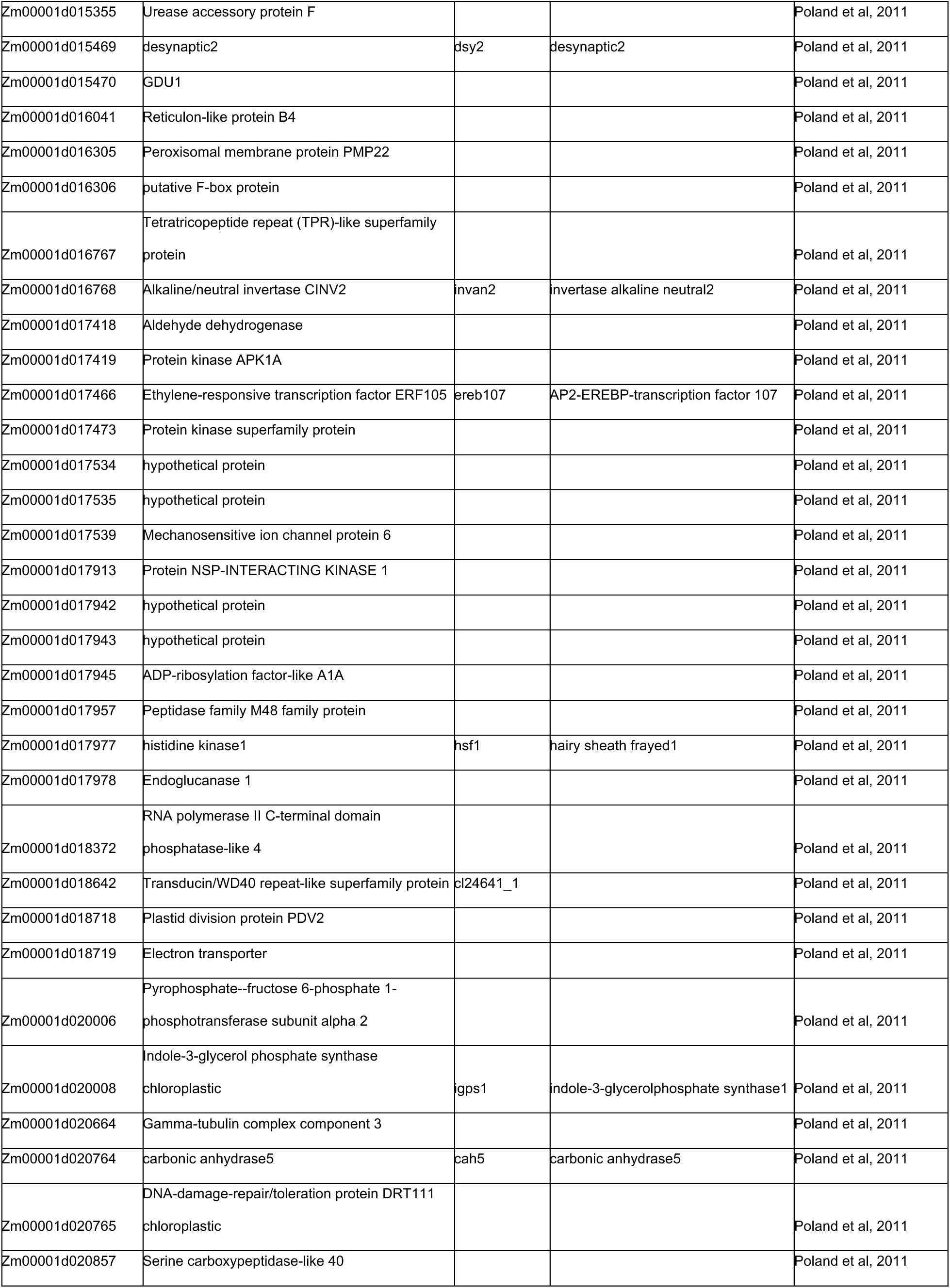
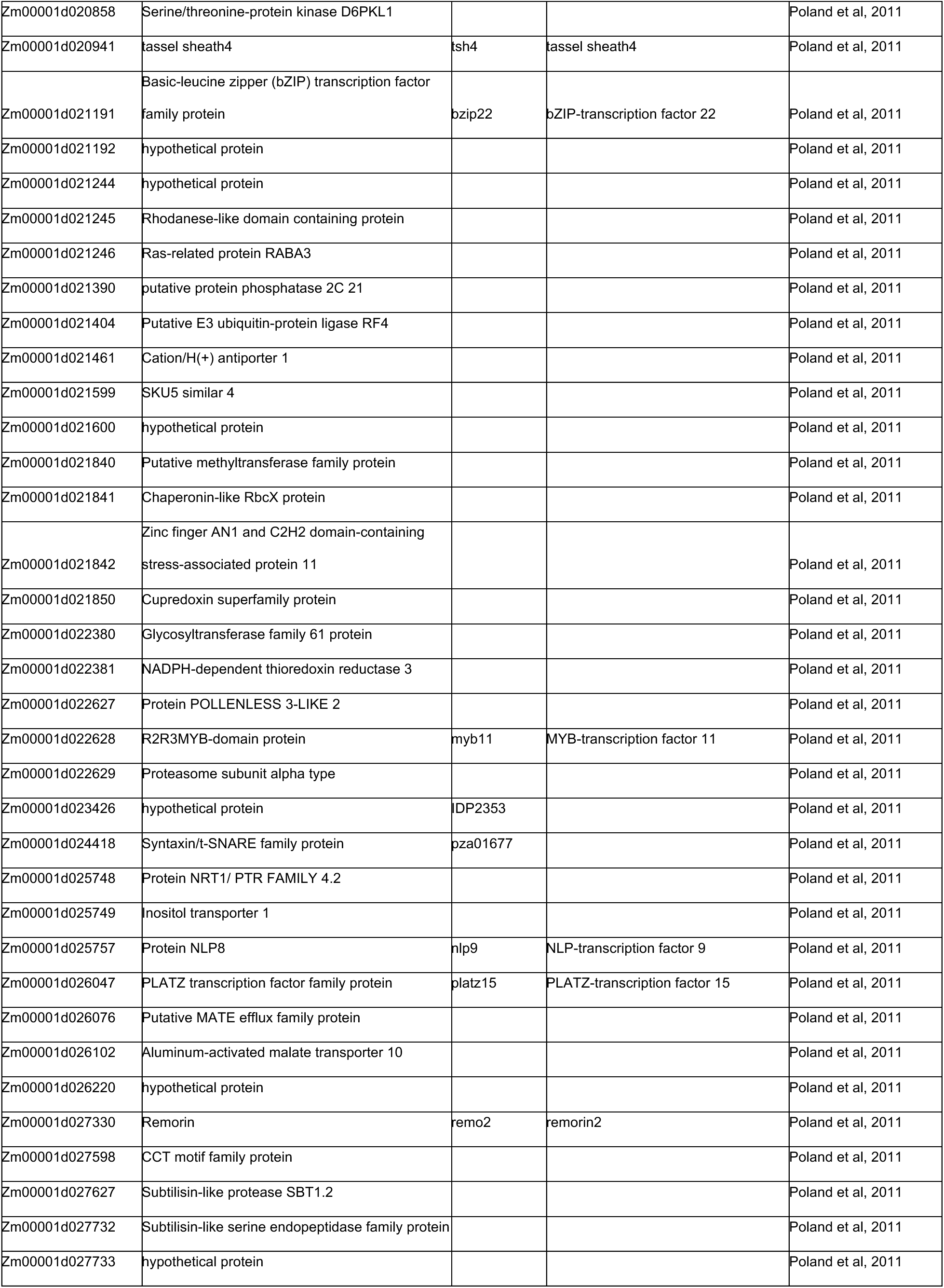
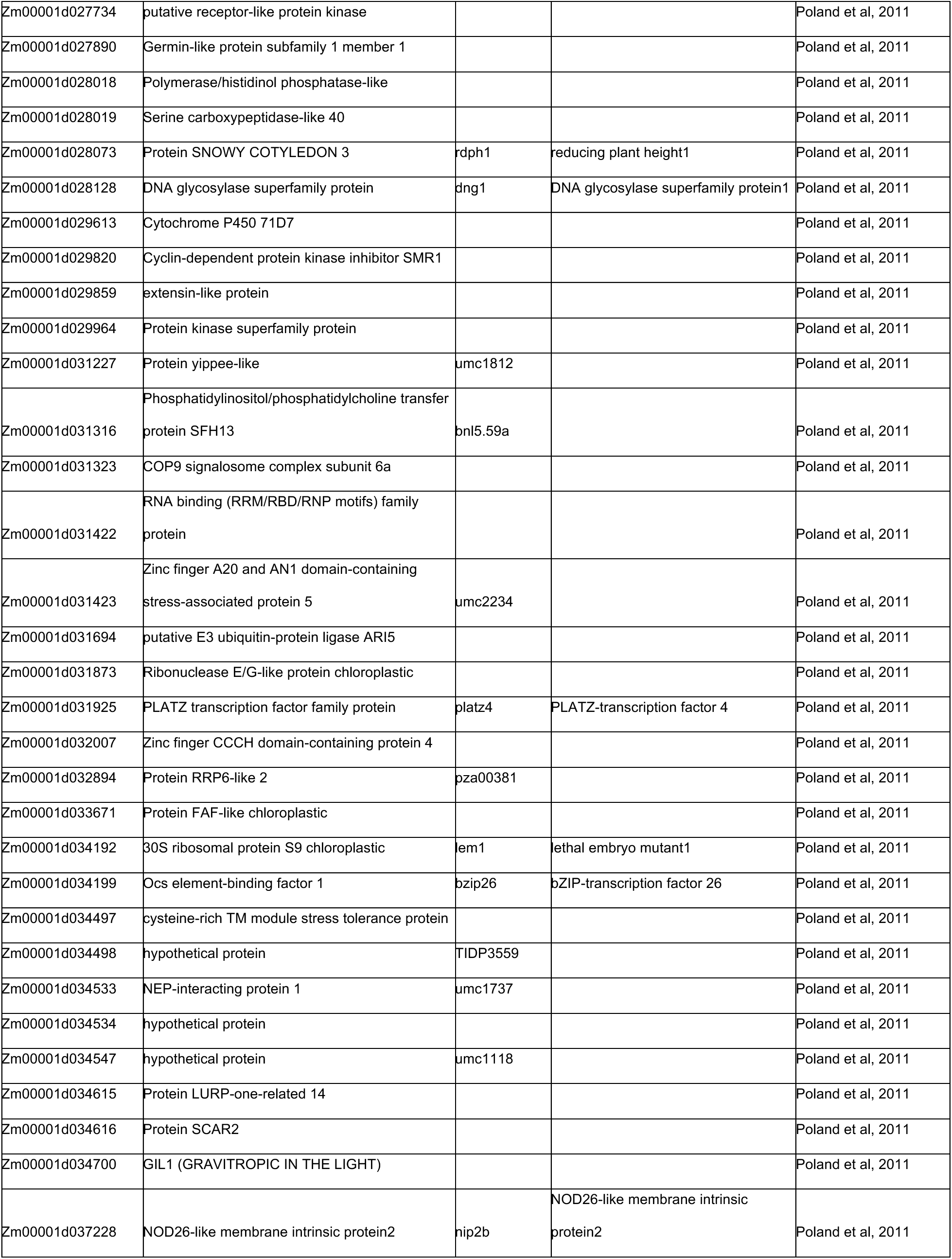
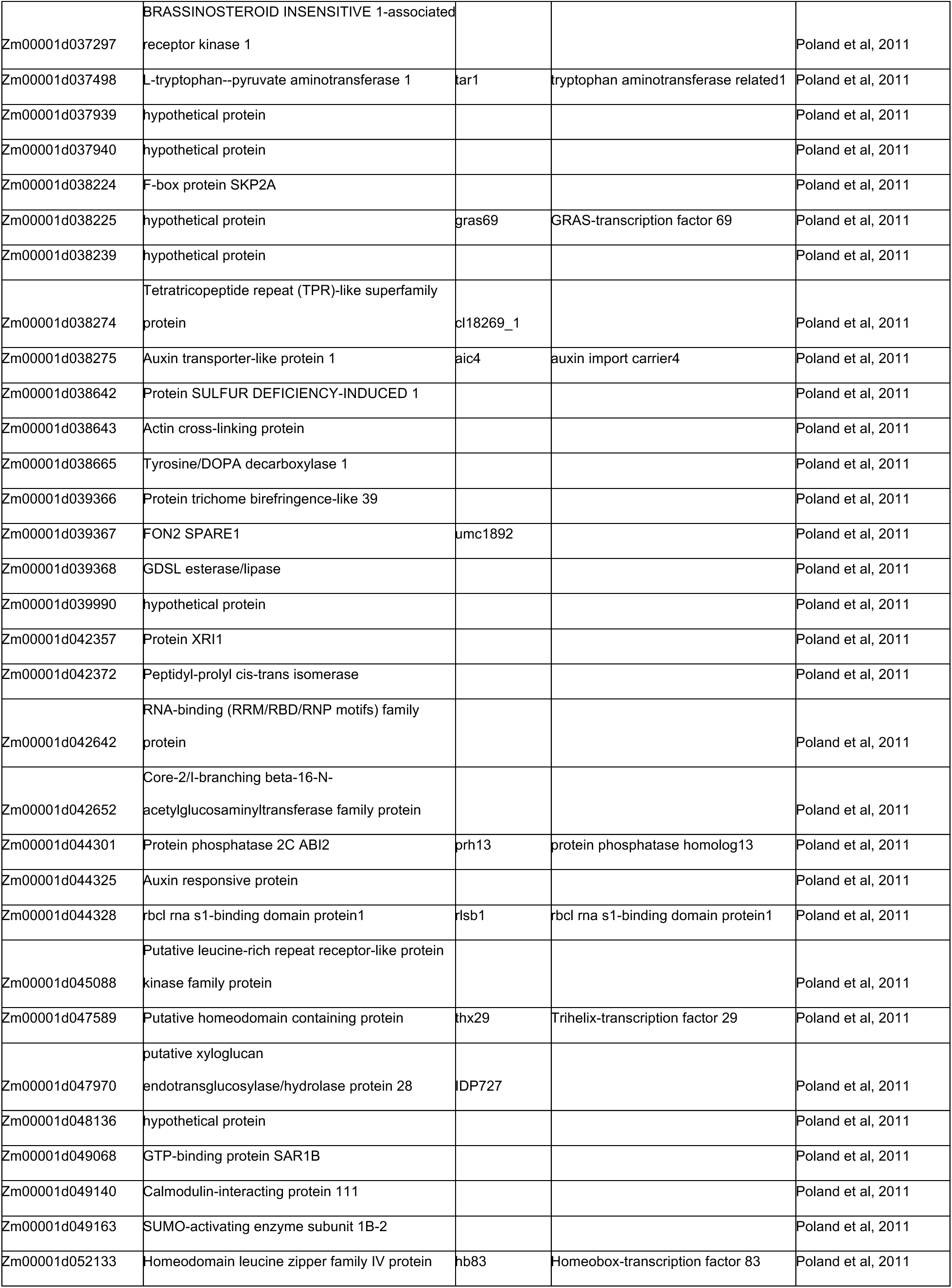
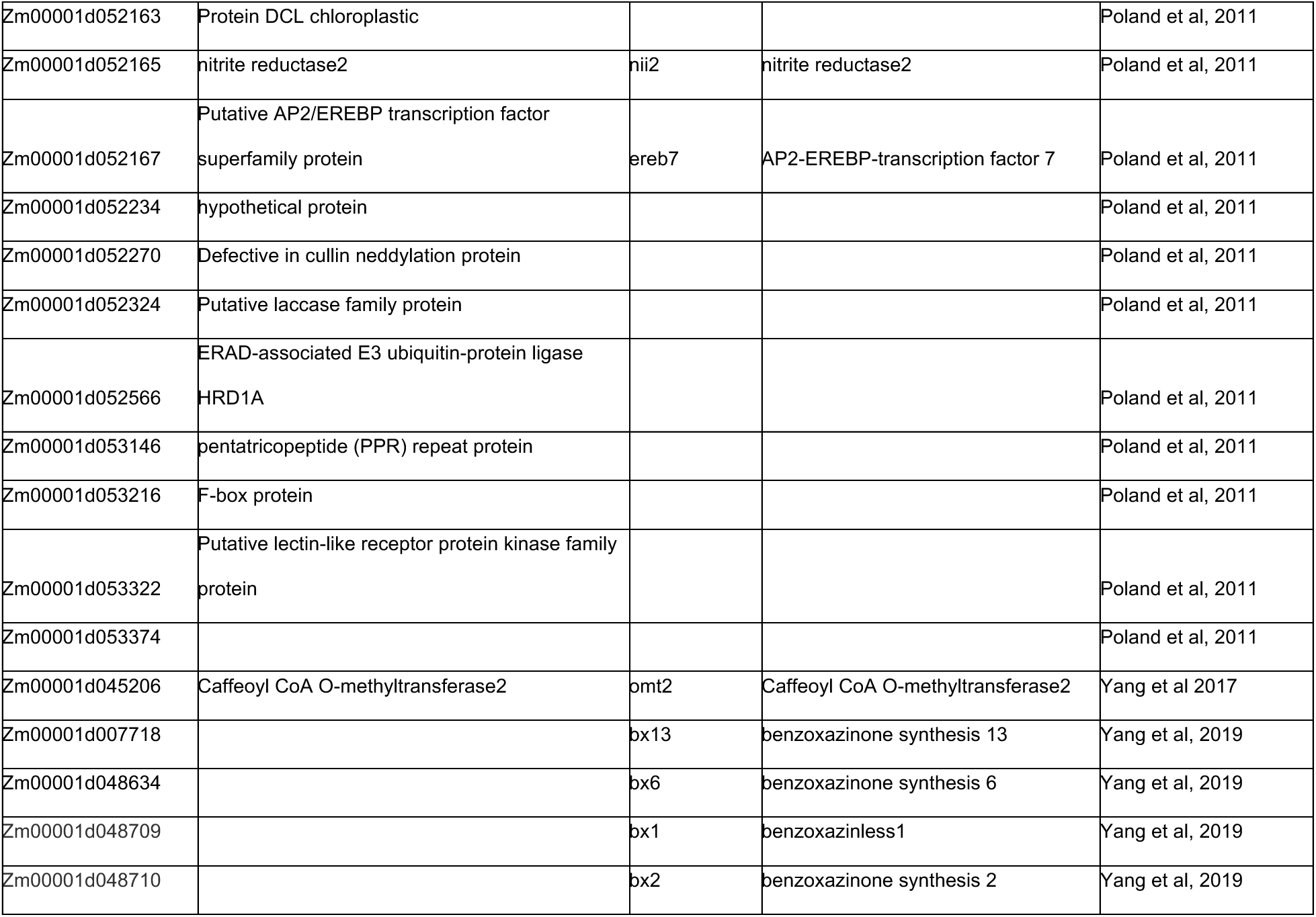
Curated list of 255 genes implicated in NLB resistance which were identified through GWAS, recombinant inbreed line (RIL) mapping, and/or mutant analysis.

**Supplemental Table 4.**
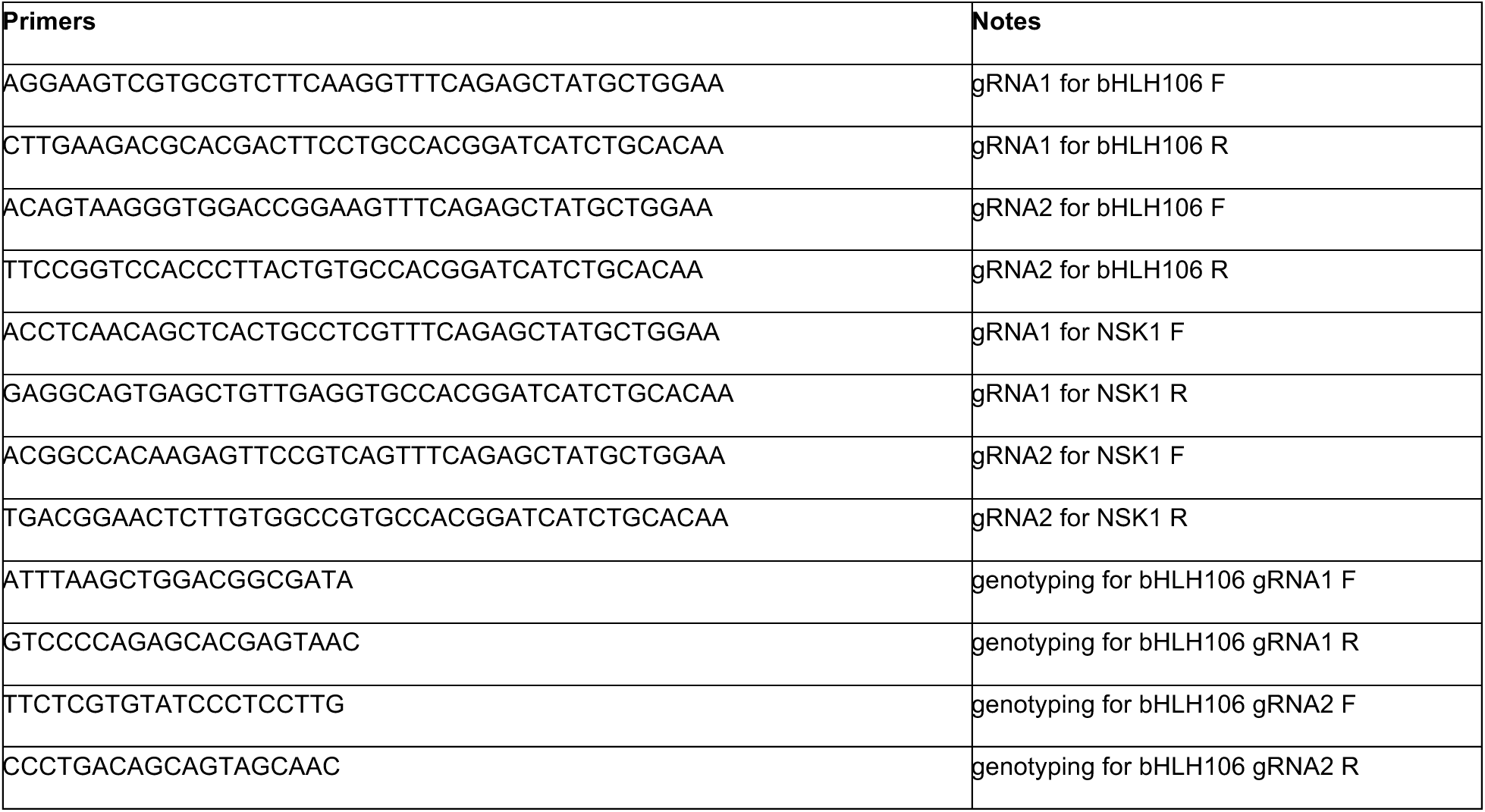
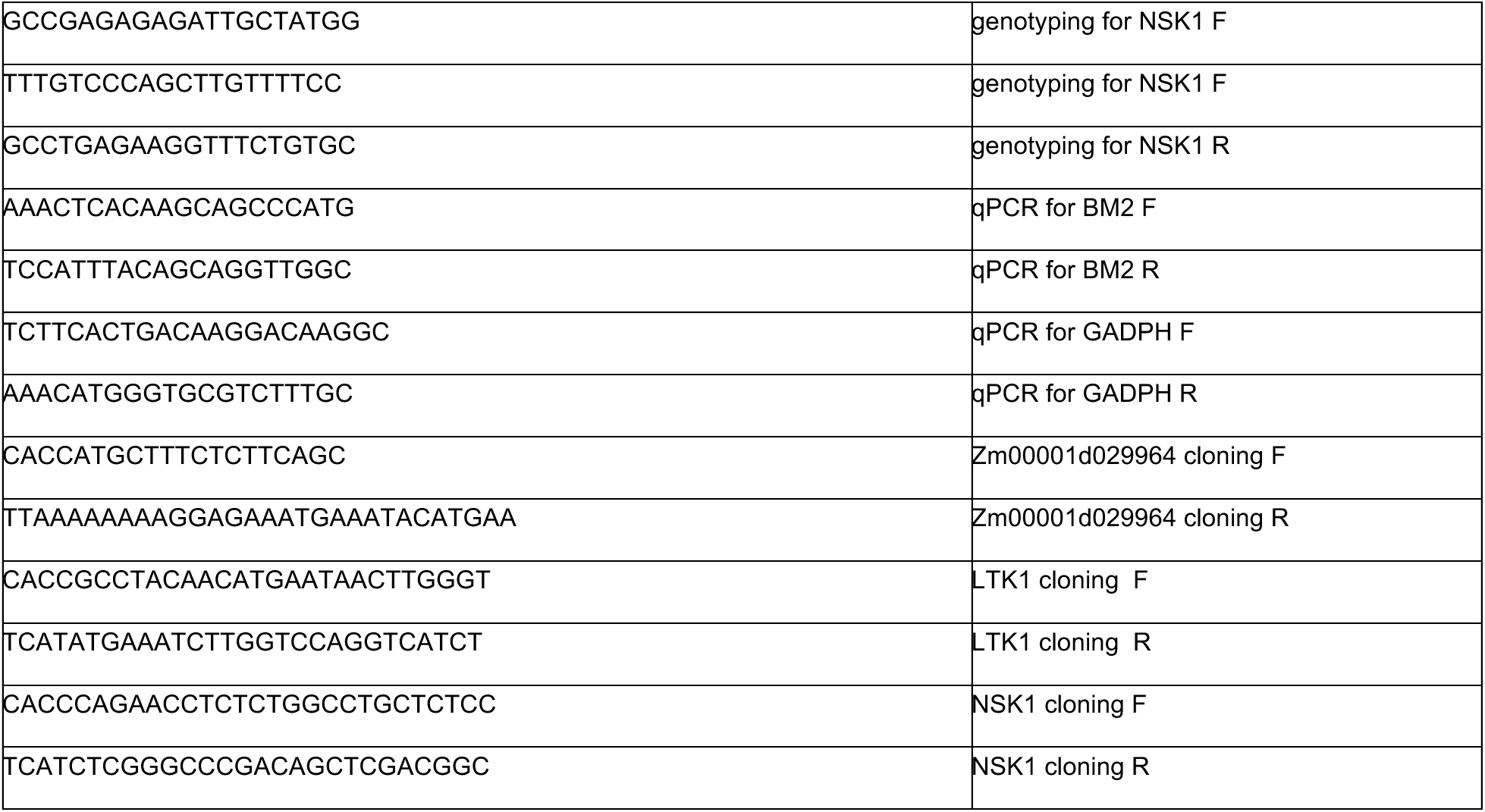
Primers used in this study.

**Supplemental Figure 1.**
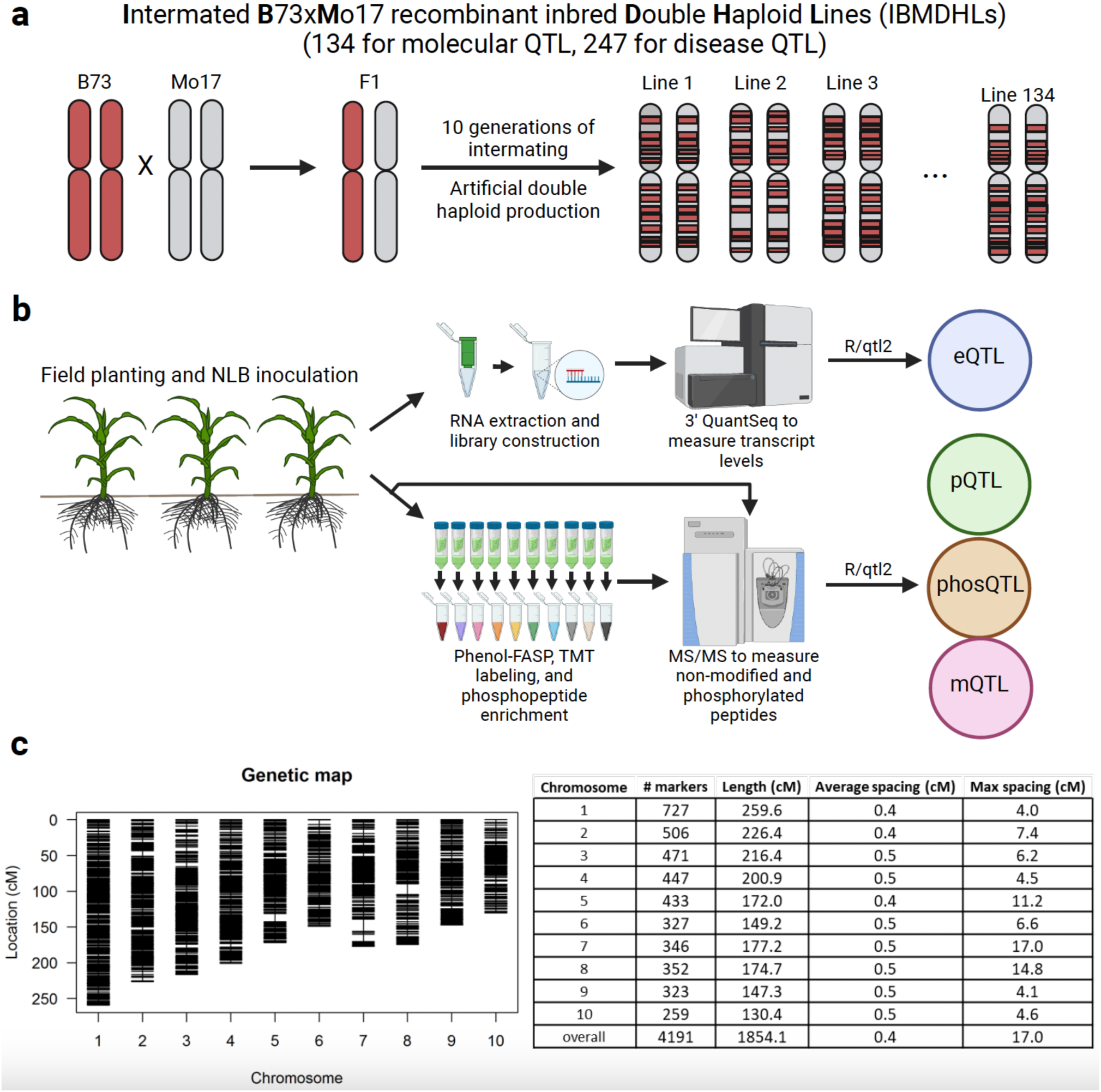
Methods for molecular and disease QTL mapping of IBMDHLs. (a) Summary of the generation of the IBMDHLs. (b) Workflow for molecular profiling of the IBMDHLs. (c) Summary of the genetic linkage map for the B73-Mo17 combined genome. (left) Diagram showing the location of the 4,191 markers (horizontal lines). (right) Coverage summary for each chromosome and overall.

**Supplemental Figure 2.**
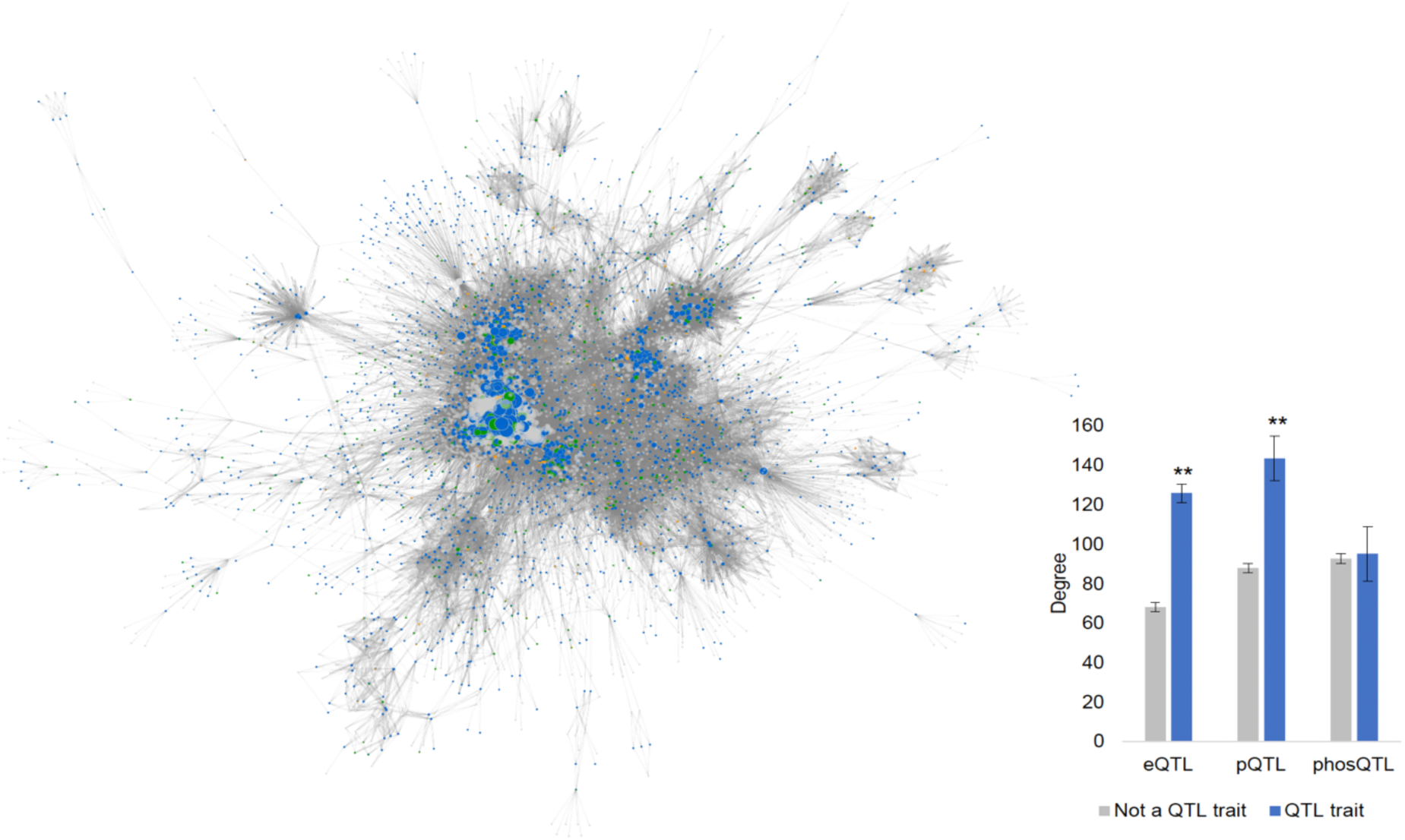
Protein-protein interaction network of molecular QTL traits. Node (protein) color represents QTL (blue: eQTL, green: pQTL, orange: phosQTL, gray: not a QTL trait). Size represents degree. (right, inset) Average degree of proteins that are a QTL trait (blue) and are not a QTL trait (gray) for each gene-product. ** denotes p < 0.0001, Wilcoxon rank-signed test.

**Supplemental Figure 3.**
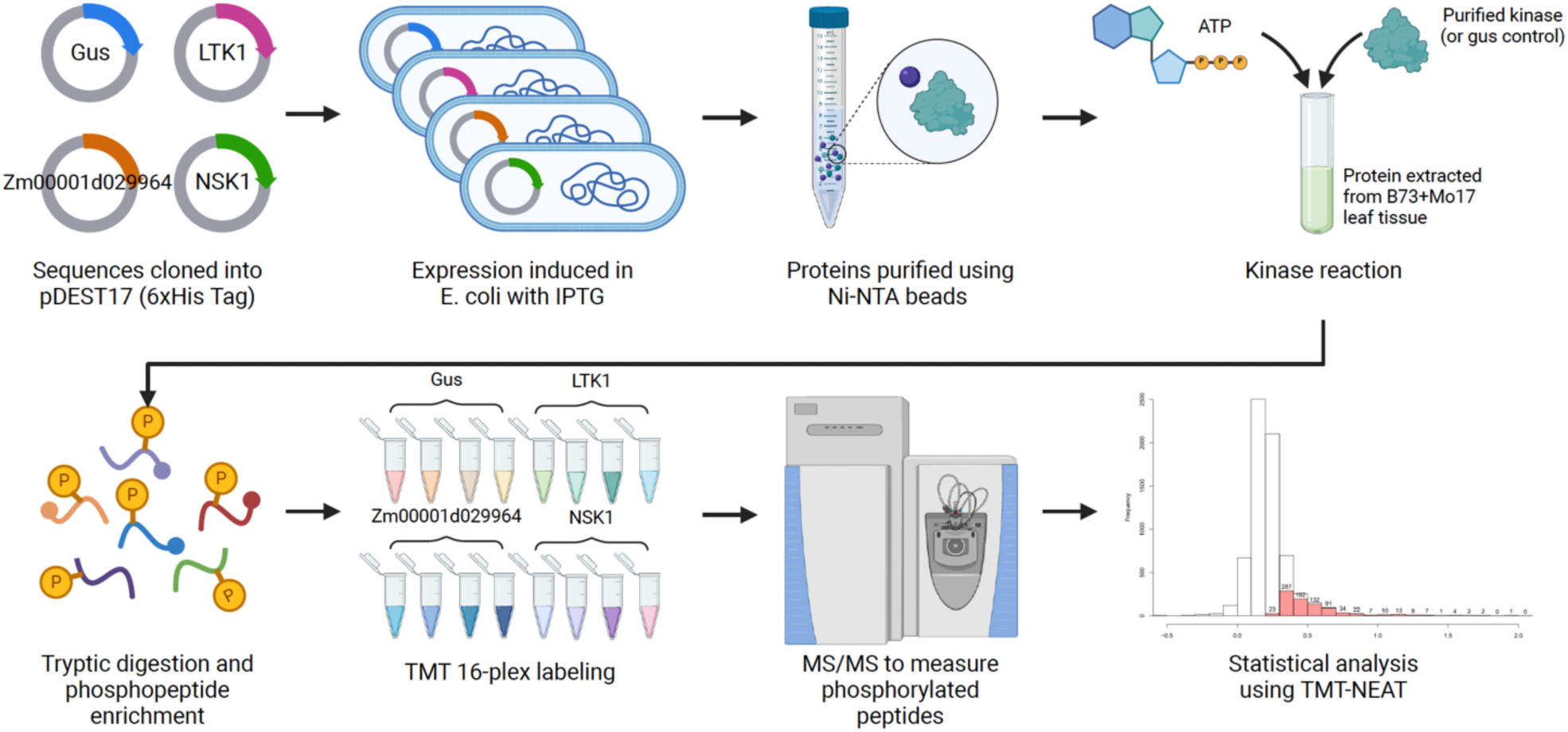
MAKS workflow to identify substrates of kinases. The kinases of interest (LTK1, Zm00001d029964, and NSK1) and the GUS control were expressed in E. coli and purified. Recombinant proteins were then incubated with proteins extracted from and equal mix of B73 and Mo17 leaf tissue. Following the kinase assays proteins were digested into peptides, TMT labeled, and phosphoproteomics was performed. Phoshosites that changed in abundance in the presence of each kinase relative to GUS were determined using TMT-NEAT.

**Supplemental Figure 4.**
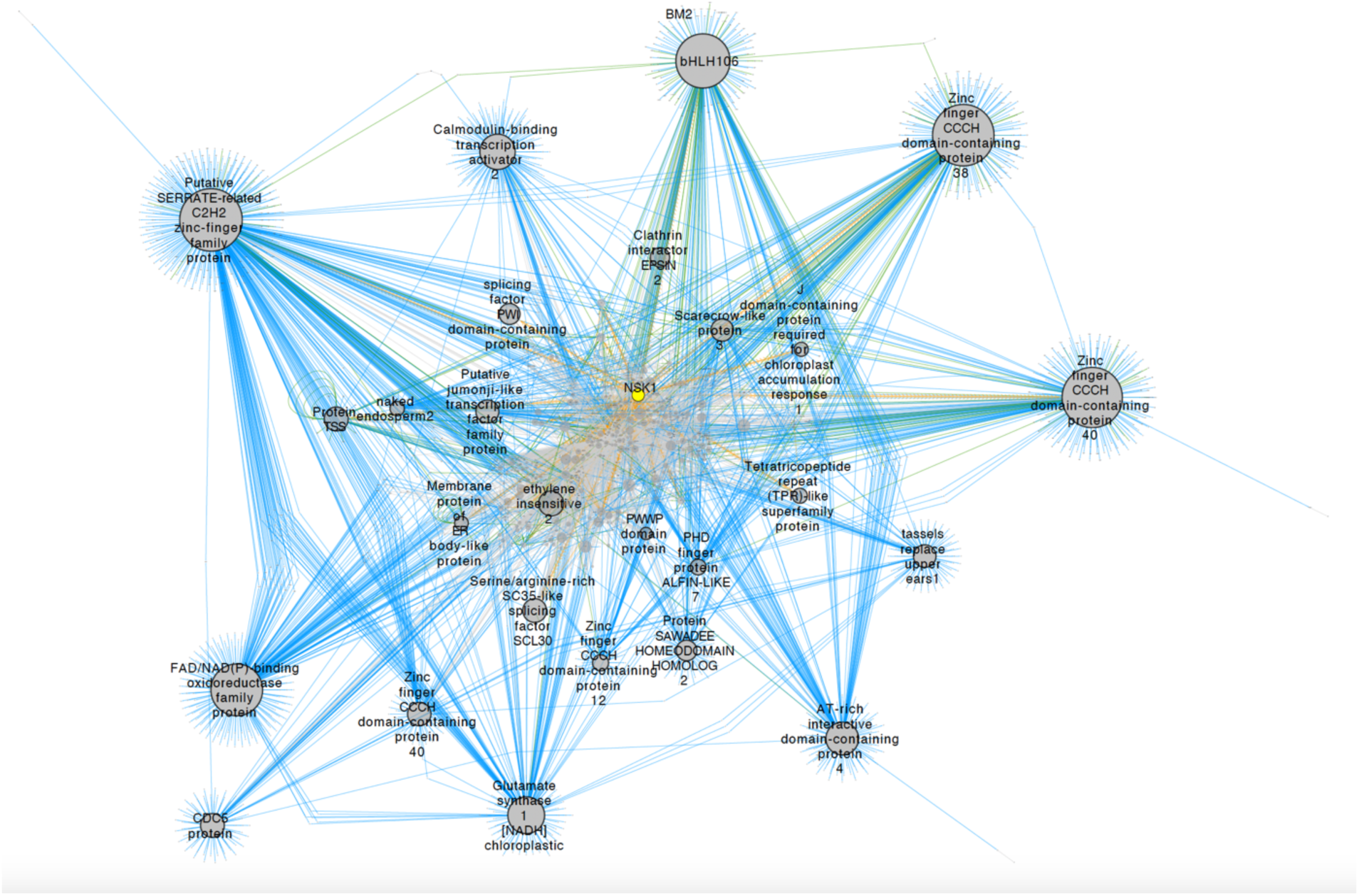
SC-ION first-neighbor network for NSK1 (yellow, center) and its downstream targets.Dark gray circles are genes. Size represents outdegree. Genes with high outdegree are outlined in black. Edge color represents QTL – eQTL (blue), pQTL (green), phosQTL (orange). Solid lines were inferred using abundance, dashed lines using phosphosite intensity, and lines with chevrons are MAKS-validated targets of NSK1. The network is arranged in an organic layout, where the top of the network is towards the center, and the bottom of the network is towards the edges.

**Supplemental Figure 5.**
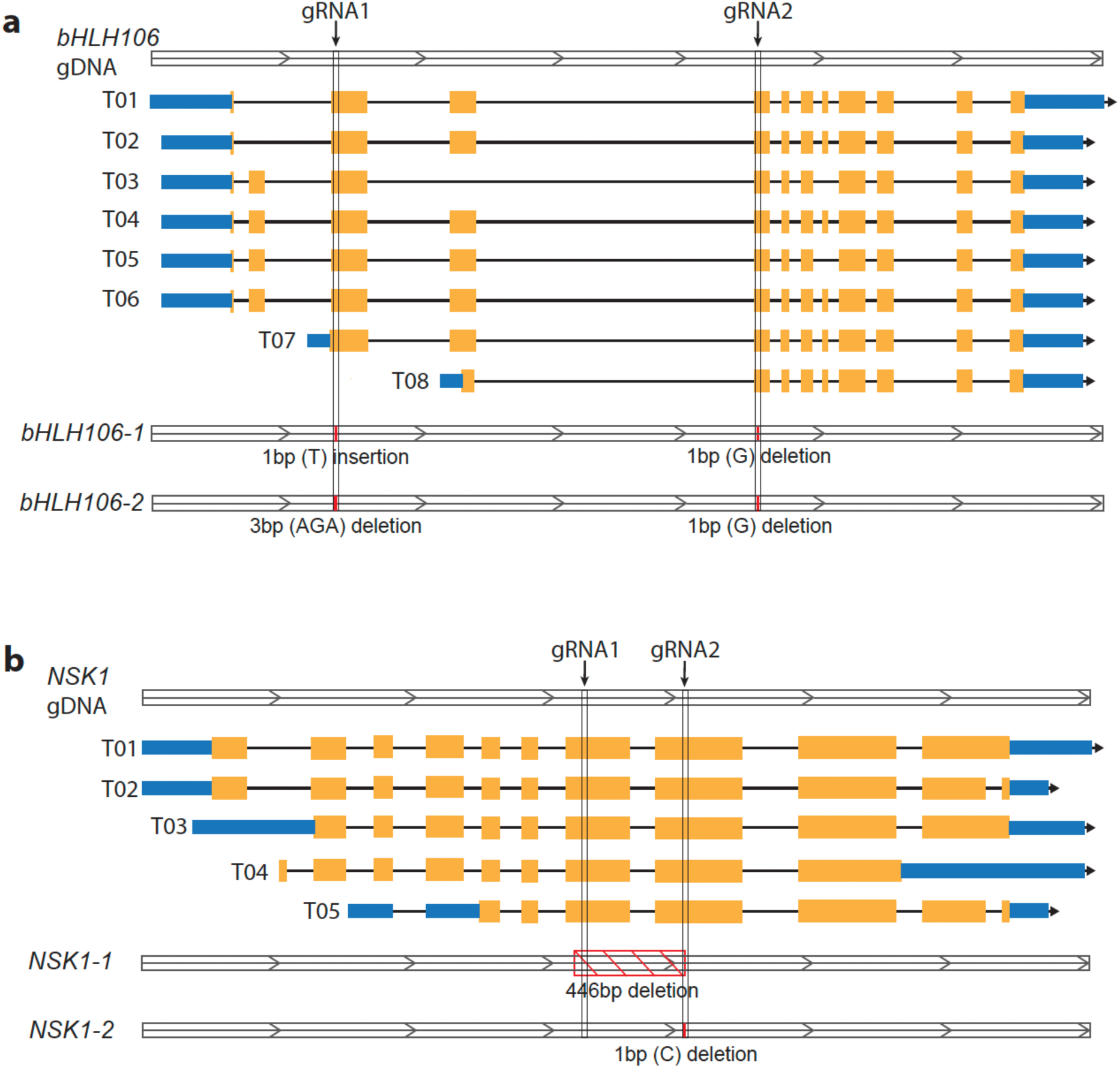
Model of CRISPR induced mutant generation for bHLH106 and NSK1. (a) Mutant generation for bHLH106 and the genotypes model of the mutant lines bHLH106-1 and bHLH106-2. (b) Mutant generation for NSK1 and the genotypes model of the mutant lines NSK1-1 and NSK1-2.The yellow fragments denotes exon region for each transcript, blue fragments denotes UTR regions and the arrow denotes the transcription direction.

## Supplemental Datasets

**Supplemental Dataset 1:** TMM-normalized expression values for all quantified transcripts.

**Supplemental Dataset 2:** Normalized protein abundance for all quantified protein groups.

**Supplemental Dataset 3:** Normalized phosphosite intensities for all quantified phosphosites.

**Supplemental Dataset 4:** B73-Mo17 genetic linkage map and genotype information.

**Supplemental Dataset 5:** All mapped molecular QTL (eQTL, pQTL, phosQTL, mQTL).

**Supplemental Dataset 6:** A list of all eQTL, pQTL, and phosQTL traits with syntenic mapping information.

**Supplemental Dataset 7:** Protein-protein interaction network from STRING database.

**Supplemental Dataset 8:** MapMan4 ontology for the B73-Mo17 combined genome and ontology enrichment analysis.

**Supplemental Dataset 9:** Metabolomics profiling results.

**Supplemental Dataset 10:** Area Under the Disease Progression Curve (AUDPC) measurements.

**Supplemental Dataset 11:** Differentially expressed gene-products in B73 and Mo17 parental lines.

**Supplemental Dataset 12:** Integrative omics networks inferred using SC-ION.

**Supplemental Dataset 13:** Multiplexed Assay for Kinase Specificity (MAKS) results.

**Supplemental Dataset 14:** A list of maize transcription factors used for SC-ION network inference.

